# Loss of the E3 ubiquitin ligases UBR-5 or HECD-1 restores *Caenorhabditis elegans* development in the absence of SWI/SNF function

**DOI:** 10.1101/2022.09.27.509717

**Authors:** Lisa Lampersberger, Francesca Conte, Subhanita Ghosh, Yutong Xiao, Jonathan Price, David Jordan, David Q Matus, Peter Sarkies, Petra Beli, Eric A Miska, Nicholas O Burton

## Abstract

SWItch/Sucrose Non-Fermenting (SWI/SNF) complexes are a family of chromatin remodellers that are conserved across eukaryotes. Mutations in subunits of SWI/SNF cause a multitude of different developmental disorders in humans, most of which have no current treatment options. Here we identify an alanine to valine causing mutation in the SWI/SNF subunit *snfc-5* (*SMARCB1* in humans) that prevents embryonic lethality in *C. elegans* nematodes harbouring a loss-of-function mutation in the SWI/SNF subunit *swsn-1* (*SMARCC1/2* in humans). Furthermore, we found that the combination of this specific mutation in *snfc-5* and a loss-of-function mutation in either of the E3 ubiquitin ligases *ubr-5* (*UBR5* in humans) or *hecd-1* (*HECTD1* in humans) can restore development to adulthood in *swsn-1* loss-of-function mutants that otherwise die as embryos. Using these mutant models, we established a set of 335 genes that are dysregulated in SWI/SNF mutants that arrest their development embryonically but exhibit near wild-type levels of expression in the presence of suppressor mutations that prevent embryonic lethality, suggesting that SWI/SNF promotes development by regulating this specific subset of genes. In addition, we show that SWI/SNF protein levels are reduced in *swsn-1; snfc-5* double mutants and partly restored to wild-type levels in *swsn-1; snfc-5; ubr-5* triple mutants, consistent with a model in which UBR-5 regulates SWI/SNF levels by tagging the complex for proteasomal degradation. Our findings establish a link between two E3 ubiquitin ligases and SWI/SNF function and suggest that UBR5 and HECTD1 might be viable therapeutic targets for the many developmental disorders caused by missense mutations in SWI/SNF subunits.

## Introduction

Chromatin remodellers are adenosine triphosphate (ATP)-powered molecular machines that can directly alter the structure of chromatin by reshuffling or evicting nucleosomes. Therefore, they control the access to DNA elements like enhancers, promoters and replication origins that need to be exposed to execute essential cellular processes such as transcription, replication and DNA repair (Saha et al., 2006). The SWItch/Sucrose Non-Fermenting (SWI/SNF) complexes were the first described chromatin remodellers, originally discovered in genetic screens in *Saccharomyces cerevisiae* in the 1980s (Neigeborn and Carlson, 1984; Stern et al., 1984). Later SWI/SNF complexes were shown to be conserved across all eukaryotes (Flaus et al., 2006). They consist of 10-15 subunits, depending on the organism (Mani et al., 2017). The human SWI/SNF complex (also known as BAF) is encoded by at least 29 genes and its core comprises one of the two mutually exclusive catalytic ATPases SMARCA2 or SMARCA4 (Kadoch and Crabtree, 2015), a hetero- or homodimer of SMARCC1/2 that acts as a scaffold for other subunits in early complex assembly (He et al., 2020; Mashtalir et al., 2018) and SMARCB1, which is important for structural complex integrity (Wang et al., 2016). Genome-wide mapping of SWI/SNF complexes by ChIP-seq and mass-spectrometry analysis of SWI/SNF co-IPs discovered diverse roles for these complexes in gene regulation and numerous interactions with other protein complexes and transcription factors (Euskirchen et al., 2011). SWI/SNF chromatin remodelling is currently estimated to regulate the expression of approximately 20% of human genes (Raab et al., 2015). This is especially important in differentiation, where SWI/SNF complexes coordinate proliferation and differentiation decisions by facilitating a balance between activation of linage-specific genes and suppression of proliferation programs (Ruijtenberg and van den Heuvel, 2016; Wilson and Roberts, 2011).

Complete loss of SWI/SNF function causes embryonic lethality in mice (Bultman et al., 2000) and even partial loss-of-function mutations in SWI/SNF chromatin remodellers cause developmental disorders such as Coffin-Siris syndrome, Nicolaides-Baraitser syndrome, Kleefstra’s syndrome, Hirschsprung’s disease, and autism in humans (Sokpor et al., 2017). The lethality observed in complete loss-of-function mutants complicates our ability to understand the mechanisms by which mutations in SWI/SNF subunits disrupt normal development (Sokpor et al., 2017). To circumvent embryonic lethality, studies have focused on studying SWI/SNF function in tissue specific mouse knock-out models (Narayanan et al., 2015) or in cell culture where individual gene knockouts are viable (Schick et al., 2019). However, cell culture models cannot recapitulate the role of SWI/SNF in animal development across tissues and the viable cell culture models are still unlikely to represent complete loss of SWI/SNF function (Schick et al., 2019).

In contrast to mammalian models, the *C. elegans* genome encodes only a single gene for each of the core SWI/SNF subunits. Deletions of the core subunits *swsn-4* (human *SMARCA2/4*) or *swsn-1* (human *SMARCC1/2*) result in embryonic or larval lethality respectively and RNAi-mediated knock-down of *snfc-5* (human *SMARCB1*) similarly results in embryonic lethality (Large and Mathies, 2014). However, previous work in *C. elegans* identified a temperature-sensitive *swsn-1* mutation. *swsn-1* temperature-sensitive mutants can develop to adulthood at the permissive temperature of 15°C but arrest in embryonic development with 100% penetrance at the restrictive temperature of 22.5°C (Sawa et al., 2000). The developmental arrest phenotype of the temperature-sensitive *swsn-1* mutants is similar to developmental defects caused by SWI/SNF mutations in humans, which suggests that SWI/SNF regulation of development is a conserved process between nematodes and humans and indicates that the temperature-sensitive *swsn-1* allele in *C. elegans* is a useful model to study SWI/SNF function in development.

Here, we report that a specific mutation in *snfc-5* (human *SMARCB1*) can prevent embryonic lethality and early developmental arrest of *swsn-1* (human *SMARCC1/2*) mutants. In addition, we report that the loss-of-function mutations in either of the genes encoding the E3 ubiquitin ligases UBR-5 or HECD-1 could further restore wild-type development in the *swsn-1* mutant model. Specifically, around 70% of hatched *swsn-1; snfc-5; ubr-5* triple mutants developed to adulthood under conditions where 100% of *swsn-1* single mutants died as embryos. Using our mutant models, we established a set of 335 genes that were specifically dysregulated in *swsn-1* mutants but exhibited near wild-type expression levels in *swsn-1; snfc-5* double and *swsn-1; snfc-5; ubr-5* triple mutants across three independent RNA-sequencing experiments, suggesting that the dysregulation of these genes drives the developmental defects observed in *swsn-1* mutants. In addition, using multiple independent approaches, we demonstrated that UBR-5 likely regulates the levels of SWI/SNF subunits to mediate its effects on SWI/SNF function. Our findings provide new insights into how defects in SWI/SNF function cause developmental defects and provide the first evidence suggesting that UBR5 or HECTD1, the human orthologs of UBR-5 and HECD-1, are potential therapeutic targets for developmental defects caused by missense mutations in SWI/SNF subunits.

## Results

### Mutations in *snfc-5*, *ubr-5*, and *hecd-1* can prevent embryonic lethality and developmental arrest in a mutant model of loss of SWI/SNF function

To identify mutations that could compensate for loss of SWI/SNF function, we utilized the *ku355* temperature-sensitive (ts) loss-of-function allele of the core SWI/SNF subunit *swsn-1* in the model animal *Caenorhabditis elegans* (Cui et al., 2004). The *swsn-1* temperature-sensitive allele encodes a P68L substitution mutation in the SWIRM protein domain of SWSN-1 (Figure S1A). 100% of animals homozygous for this mutation die as embryos when grown at 22.5°C or arrest at early larval stages of development when exposed to high temperatures after completing embryonic development due to a lack of SWI/SNF function (Sawa et al., 2000). We mutagenized *swsn-1* mutants with ethyl methanesulfonate (EMS) at a permissive temperature (20°C) and subjected their F3 offspring as synchronized embryos to the restrictive temperature (25°C) for 72 hours (Figure 1SB). We identified five mutant isolates that did not arrest development at early larval stages. By performing whole genome sequencing of these five isolates, we found that one of the recovered isolates carries an additional *swsn-1* substitution mutation (V62I) nearby the original P68L mutation. This mutation is likely an internal suppressor and was not further validated. The remaining four isolates from this screen, which were all from independent pools, all carry an identical A258V substitution mutation in the gene encoding SNFC-5 (Figure S1C), another core subunit of the SWI/SNF complex and homolog of human SMARCB1. We recreated the A258V *snfc-5* mutation by CRISPR-Cas9 gene editing (Paix et al., 2015) and confirmed that this mutation prevents early larval arrest in *swsn-1* mutants (Figure S1D-F). We conclude that the A258V mutation in *snfc-5* can suppress some of the developmental defects observed in *swsn-1* mutants.

We found that exposure of mutant L4-staged or young adult animals to 25°C for 16 hours prior to collecting the embryos resulted in 100% embryonic lethality for *swsn-1* single mutants and 95% of *swsn-1; snfc-5* double mutants (Figure S1G). Of the few surviving *swsn-1; snfc-5* double mutants 98% arrested development between L1 and L3 stages (Figure 1A-B and S1H). These findings indicate that this specific mutation in *snfc-5* can suppress the embryonic lethality in a proportion of *swsn-1* mutants but is not sufficient to restore development to adulthood in the animals that do not die as embryos. Therefore, we asked whether we could suppress the developmental defects of *swsn-1; snfc-5* double mutants further by introducing additional mutations and if this would allow us to identify novel genetic interactors of the SWI/SNF complex. To test this hypothesis, we performed a second EMS mutagenesis screen with *swsn-1; snfc-5* double mutants using these more stringent conditions and screened for mutants that could develop to adulthood (Figure S1I). From this screen we identified and sequenced the genomes of 12 independently isolated mutants that developed to adulthood. We found that two of the mutant isolates have additional mutations in *swsn-1* or *snfc-5* (Figure S1J). These mutations are likely internal suppressor mutations and were not characterized further. Seven of the remaining isolates each carry different predicted loss-of-function mutations in the gene encoding the HECT-type E3 ubiquitin ligase UBR-5 and three have mutations in the HECT-type E3 ubiquitin ligase HECD-1, a paralog of UBR-5 (Figure 1C and S1J). We recreated the UBR-5 Q150* allele, as it was the earliest pre-mature stop mutation we identified (see Figure 1C), by CRISPR-Cas9 gene editing (Paix et al., 2015) (*mj638*). We then crossed *swsn-1; snfc-5* double mutants to *ubr-5(mj638)* mutants and confirmed that this new mutation in *ubr-5* restored development to at least the L4 larval stage in approximately 70% of viable animals in the SWI/SNF mutant model (Figure 1A-B and S1H). Moreover, we found that approximately 29% of *swsn-1; snfc-5; ubr-5* triple mutants did not die as embryos, which is a more than 5-fold decrease of embryonic lethality compared to *swsn-1; snfc-5* double mutants (Figure S1G). Similarly, we crossed *swsn-1; snfc-5* double mutants with mutants harbouring a deletion in *hecd-1* (*ok1437)* and again confirmed that the loss of HECD-1 (introduction of the predicted *hecd-1* null allele) restored development to the L4 larval stage in 46% of animals in our SWI/SNF mutant model (Figure 1A-B and S1H). We conclude that mutations in *ubr-5* and *hecd-1* can restore development to adulthood in *swsn-1; snfc-5* double mutant animals.

**Figure 1:**
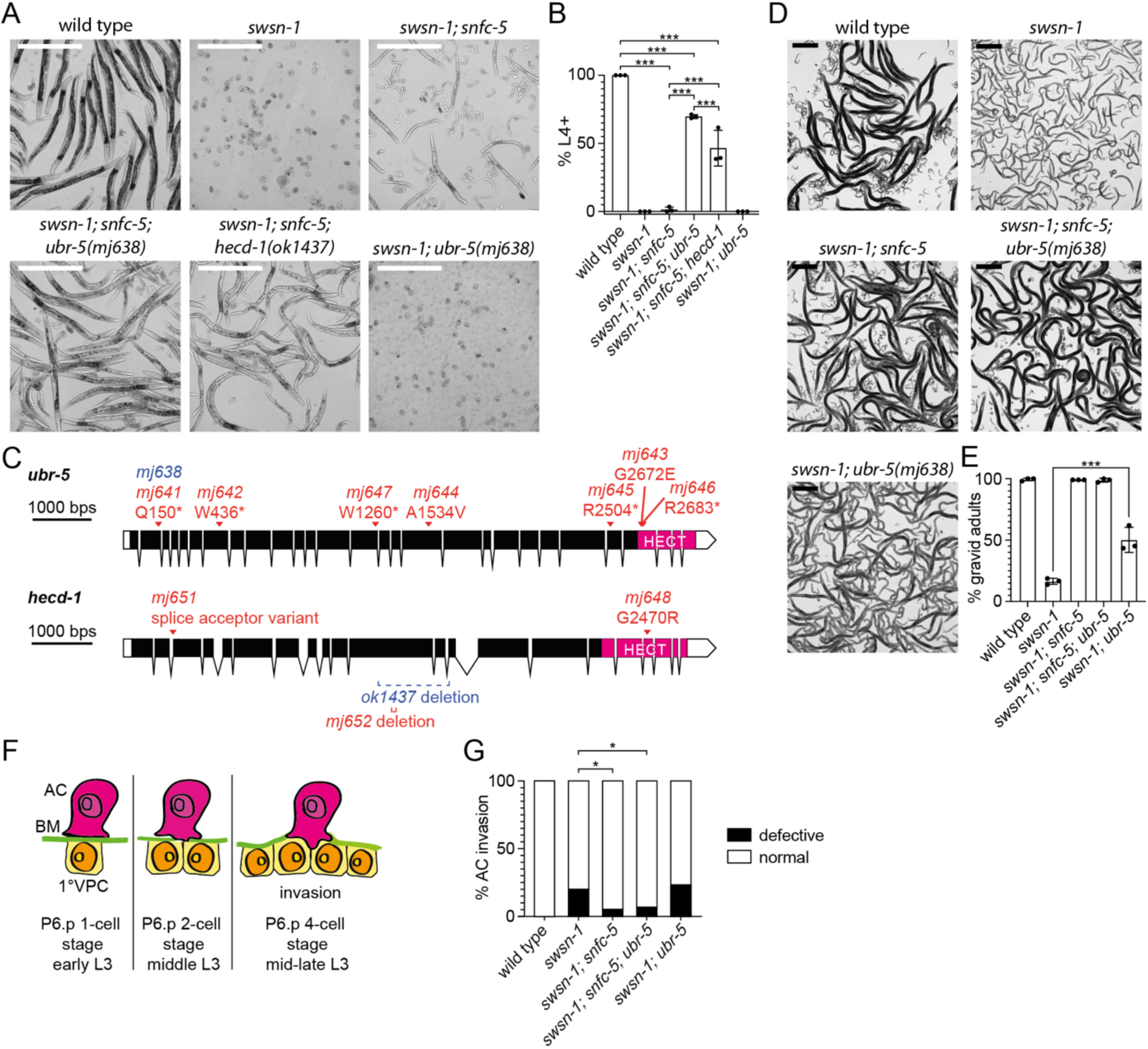
Mutations in *snfc-5*, *ubr-5*, and *hecd-1* can prevent embryonic lethality and developmental arrest of *swsn-1* mutants. **A-B)** Quantification of *C. elegans* developmental stages after exposing the parental generation to 25°C for 16 hours and collecting and growing embryos at 25°C for 72 hours. (A) Representative images of wild-type animals, *swsn-1* single, *swsn-1; snfc-5* double, *swsn-1; snfc-5; ubr-5* triple, *swsn-1; snfc-5; hecd-1* triple and *swsn-1; ubr-5* double mutants, scale bar = 500μm. (B) Percentage of L4 stage or older animals (n=3 of >= 100 animals), bar heights represent the mean, error bars represent standard deviation, *** = Bonferroni corrected Fisher’s exact test p-value < 0.0001, calculated using contingency table in Figure S1H. **C)** Schematic representation of *ubr-5* and *hecd-1* with alleles identified in the second EMS screen (red), CRISPR recreated Q150* *ubr-5* allele (blue) and available *hecd-1* deletion allele *ok1437* (blue). C-terminal HECT domains are indicated in pink. Graphic made using http://wormweb.org/exonintron. **D-E)** Quantification of *C. elegans* developmental stages after embryos were exposed to 22.5°C for 48 hours and 25°C for 24 hours. (D) Representative images of wild-type animals, *swsn-1* single, *swsn-1; snfc-5;* double, *swsn-1; snfc-5; ubr-5* triple and *swsn-1; ubr-5* double mutants, larvae in the images of the wild-type, *swsn-1; snfc-5* double and *swsn-1; snfc-1; ubr-5* triple mutants are the offspring of scored animals and were not scored, scale bar = 500μm. (E) Percentage of gravid adults (n=3 of >100 scored animals), bar heights represent the mean, error bars represent standard deviation, *** = Bonferroni corrected Fisher’s exact test p-value < 0.0001, calculated using contingency table in Figure S1K. **F-G)** Quantification of anchor cell (AC) invasion. (F) Schematic of AC invasion, BM = basement membrane, 1° VPC = primary vulval precursor cell. (G) Scoring of invasion defects, * = Fisher’s exact test p-value < 0.05, calculated using contingency table in Figure S1M. Alleles used: *swsn-1(ku355), snfc-5(mj633), ubr-5(mj638), hecd-1(ok1437)*.

To test if the loss of UBR-5 (introduction of the predicted *ubr-5* null allele) is sufficient to restore developmental defects in *swsn-1* single mutants even in the absence of the *snfc-5* mutation, we generated *swsn-1; ubr-5* double mutants and assayed their development under different conditions. We found that loss of *ubr-5* alone increased the number of *swsn-1* mutants that developed into gravid adults from 14% to 45% when embryos were grown at room temperature for 48 hours and then shifted to 25°C for 24 hours (Figure 1D-E and S1K) but was not sufficient to restore developmental defects under more stringent conditions in which *swsn-1* single mutants died as embryos (Figure 1A-B and S1H). These findings indicate that the mutation in *snfc-5* is required to suppress the embryonic lethality caused by mutation in *swsn-1*, but that the loss of UBR-5 is sufficient to improve development from larvae to gravid adults in *swsn-1* mutants under less stringent conditions.

Mutations in SWI/SNF subunits cause numerous defects in animal development and physiology. For example, the *swsn-1* mutation also causes defective anchor cell (AC) invasion, a process required for establishing the uterine-vulval connection during larval development critical for adult egg-laying (Smith et al., 2022) (see schematic of AC invasion in Figure 1F). To test whether the suppressor mutations we identified specifically suppress the developmental arrest in *swsn-1* mutants or if they might generally suppress the loss of SWI/SNF function, we assayed AC invasion in wild-type, *swsn-1* mutants, and various combinations of double and triple mutant animals. We found that *swsn-1* single mutants exhibited defective AC invasion phenotype in 20% (10/50 animals) of animals (Figure 1G and S1L-M), consistent with previously published findings (Smith et al., 2022). This phenotype was partially rescued in *swsn-1; snfc-5* double (5.2% invasion defects, 4/77 animals) and *swsn-1; snfc-5; ubr-5* triple mutants (6.78% invasion defects, 5/74 animals), but not in *swsn-1; ubr-5* double mutants (23% invasion defects, 7/30 animals) (Figure 1G and S1L-M). These data suggest that the mutation in *snfc-5* generally suppresses many of the defects observed in *swsn-1* single mutants, but that the loss of UBR-5 specifically suppresses the developmental arrest caused by loss of SWI/SNF function.

### Loss of the UBR-5 key catalytic residue is sufficient to suppress the developmental arrest of SWI/SNF mutants

HECT-type E3 ubiquitin ligases such as UBR-5 and HECD-1 have a catalytic cysteine within their C-terminal HECT domains (Wang et al., 2020). Ligation reactions depend on this catalytic cysteine, which forms a thioester-linked intermediate with the ubiquitin before ligating it onto a substrate protein (Wang et al., 2020). We replaced the catalytic residue of UBR-5 (cysteine 2913) with an alanine or serine by CRISPR-Cas9 gene editing (Paix et al., 2015) (Figure 2A). The C2913A substitution mutation should prevent the loading of ubiquitin onto UBR-5, whereas the C2913S substitution mutation should still enable the loading, but prevent the transfer of ubiquitin onto a substrate (Garcia-Barcena et al., 2020). We found that substitution of the catalytic cysteine of UBR-5 to alanine or serine in *swsn-1; snfc-5* double mutants resulted in a similar suppression of the temperature-sensitive larval arrest as introducing the premature stop *ubr-5* allele (*mj638*) (Figure 2B-C and S2). These results indicate that inactivation of the catalytic function of UBR-5 is sufficient for the suppression of developmental arrest of *swsn-1; snfc-5* mutants.

**Figure 2:**
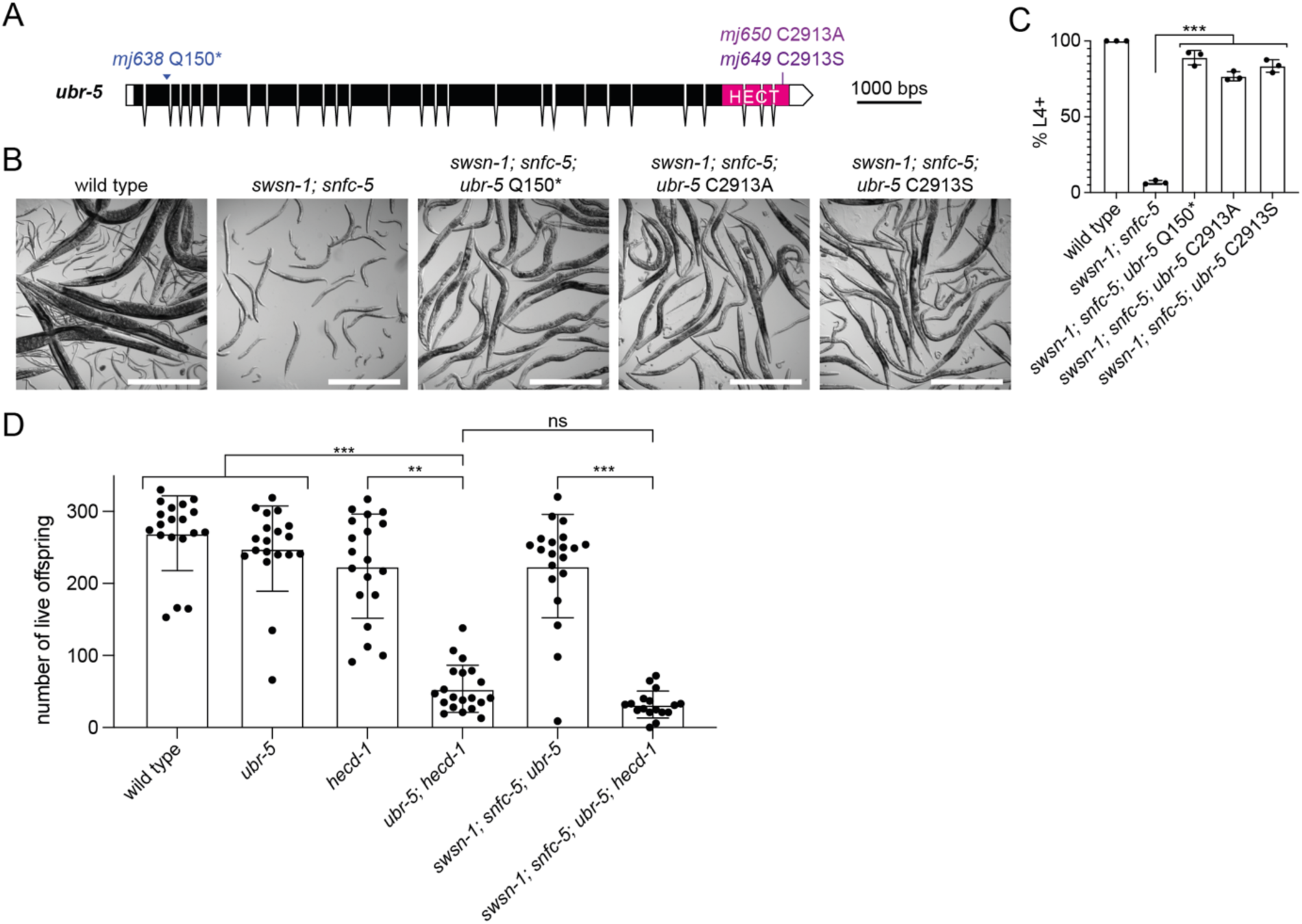
Loss of key catalytic residue of UBR-5 is sufficient for developmental arrest suppression and the combined loss of UBR-5 and HECD-1 have deleterious effects on *C. elegans* reproduction. **A)** Schematic representation of *ubr-5* with CRISPR generated Q150*, C2913A and C2913S substitutions. C-terminal HECT protein domains are indicated in pink. **B-C)** Quantification of *C. elegans* developmental stages after exposing the parental generation to 25°C for 16 hours and collecting and growing embryos at 25°C for 72 hours. (B) Representative images of wild-type animals (larvae in this image are the offspring of scored animals and were not scored), *swsn-1; snfc-5* double, *swsn-1; snfc-5*; *ubr-5* Q150* triple, *swsn-1; snfc-5; ubr-5* C2913A triple and *swsn-1; snfc-5; ubr-5* C2913S triple mutants, scale bar = 500μm. (C) Percentage of animals L4 larval stage or older (n=3 of >=100 scored animals), bar heights represent the mean, error bars represent standard deviation, *** = Bonferroni corrected Fisher’s exact test p-value < 0.0001, calculated using contingency table in Figure S2. **D)** Number of live offspring of individual wild-type animals, *ubr-5* single, *hecd-1* single, *ubr-5; hecd-1* double, *swsn-1; snfc-5; ubr-5* triple and *swsn-1; snfc-5; ubr-5; hecd-1* quadruple mutant animals, bar heights represent the mean and error bars represent standard deviation, Kruskal-Wallis test (H_5_ = 74.76, p < 0.001), Dunn’s multiple comparison adjusted p- value ** = < 0.001, *** = < 0.0001, ns = not significant. Alleles used: *swsn-1(ku355), snfc-5(mj633), ubr-5(mj638)* (this *ubr-5* allele was used in D)*, ubr-5(mj650), ubr-5(mj649), hecd-1(ok1437)*.

### The combined loss of UBR-5 and HECD-1 has deleterious effects on *C. elegans* reproduction

Since the loss of either UBR-5 or HECD-1 restored development to adulthood in some of the *swsn-1; snfc-5* double mutants (Figure 1A-B and S1H), we wondered whether the combined loss of both E3 ubiquitin ligases would have an even greater effect. However, when generating *swsn-1; snfc-5; ubr-5; hecd-1* quadruple mutants, we observed that those animals were substantially sicker and had fewer offspring than other mutant combinations. This effect appears to be a synthetic interaction between *ubr-5* and *hecd-1* because we found that *ubr-5; hecd-1* double mutants exhibited a similar phenotype even in the absence of any SWI/SNF subunit mutations. Specifically, we found that the loss of either of the paralogous HECT-type E3 ubiquitin ligases alone did not affect animal reproduction, but the loss of both UBR-5 and HECD-1 resulted in a substantial reduction of live offspring. On average the *ubr-5; hecd-1* double mutants had about four-times fewer live offspring than either of the *ubr-5* or *hecd-1* single mutants (Figure 2D). The quadruple mutants also had significantly fewer live offspring compared to the *swsn-1; snfc-5; ubr-5* triple mutants and did not have significantly fewer offspring compared to the *ubr-5; hecd-1* double mutants (Figure 2D). Thus UBR-5 and HECD-1 likely have redundant functions and loss of one can be compensated by the other protein. To our knowledge, these findings are the first to indicate that these two ubiquitin ligases might function redundantly.

### UBR-5 regulates SWI/SNF protein levels

Our findings suggest that the catalytic activity of UBR-5 is involved in supressing the *swsn-1* mutant developmental arrest (Figure 2B-C and S2). One of the best understood roles of ubiquitin ligation to proteins is to tag them for proteasomal degradation (Akutsu et al., 2016). The human homologs of UBR-5 and HECD-1, UBR5 and HECTD1 respectively, have both been shown to mediate K48-linked ubiquitination (Wang et al., 2020), which is a signal for proteasomal degradation (Zheng and Shabek, 2017). A possible and direct link between the SWI/SNF complex and these two ubiquitin ligases could be that the *swsn-1* mutation destabilizes the SWI/SNF complex and that these enzymes ubiquitinate unstable complexes to promote their degradation. In this case, loss of UBR-5 should result in higher levels of SWSN-1 which in turn might explain the observed developmental arrest suppression. To test this hypothesis, we measured SWSN-1 protein levels in wild-type animals, *swsn-1* single, *swsn-1; snfc-5* double and *swsn-1; snfc-5; ubr-5* triple mutants by Western blotting. We generated endogenously FLAG-tagged wild-type and mutant versions of SWSN-1 (Figure 3A). We synchronized L1-staged animals by starvation and subsequently fed them and exposed them to the restrictive temperature (25°C) for six hours. These conditions were chosen so that all of the mutants would be at closely matched developmental stages. We found that levels of SWSN-1 were on average approximately 40% reduced in *swsn-1* single mutants when compared to wild-type animals, suggesting that the P86L mutation reduces SWSN-1 protein levels. This reduction in SWSN-1 levels was largely restored in *swsn-1; snfc-5; ubr-5* triple mutants (Figure 3B). However, these effects were highly variable and the levels of SWSN-1 were not statistically significantly different between *swsn-1; snfc-5* double mutants and *swsn-1; snfc-5; ubr-5* triple mutants (Figure 3B) even though we found that most *swsn-1; snfc-5* double mutants arrested their development at early larval stages while most *swsn-1; snfc-5; ubr-5* triple mutants were able to develop to L4-stages and adulthood (Figure 1A-B and S1H). These results suggest that mutation of *swsn-1* leads to a reduction of SWSN-1 protein levels. Furthermore, UBR-5 potentially has either a small effect on SWSN-1 protein levels or an effect that occurs only in specific cells or at specific developmental stages that is difficult to detect when measuring SWSN-1 levels in whole animals by Western blotting.

**Figure 3:**
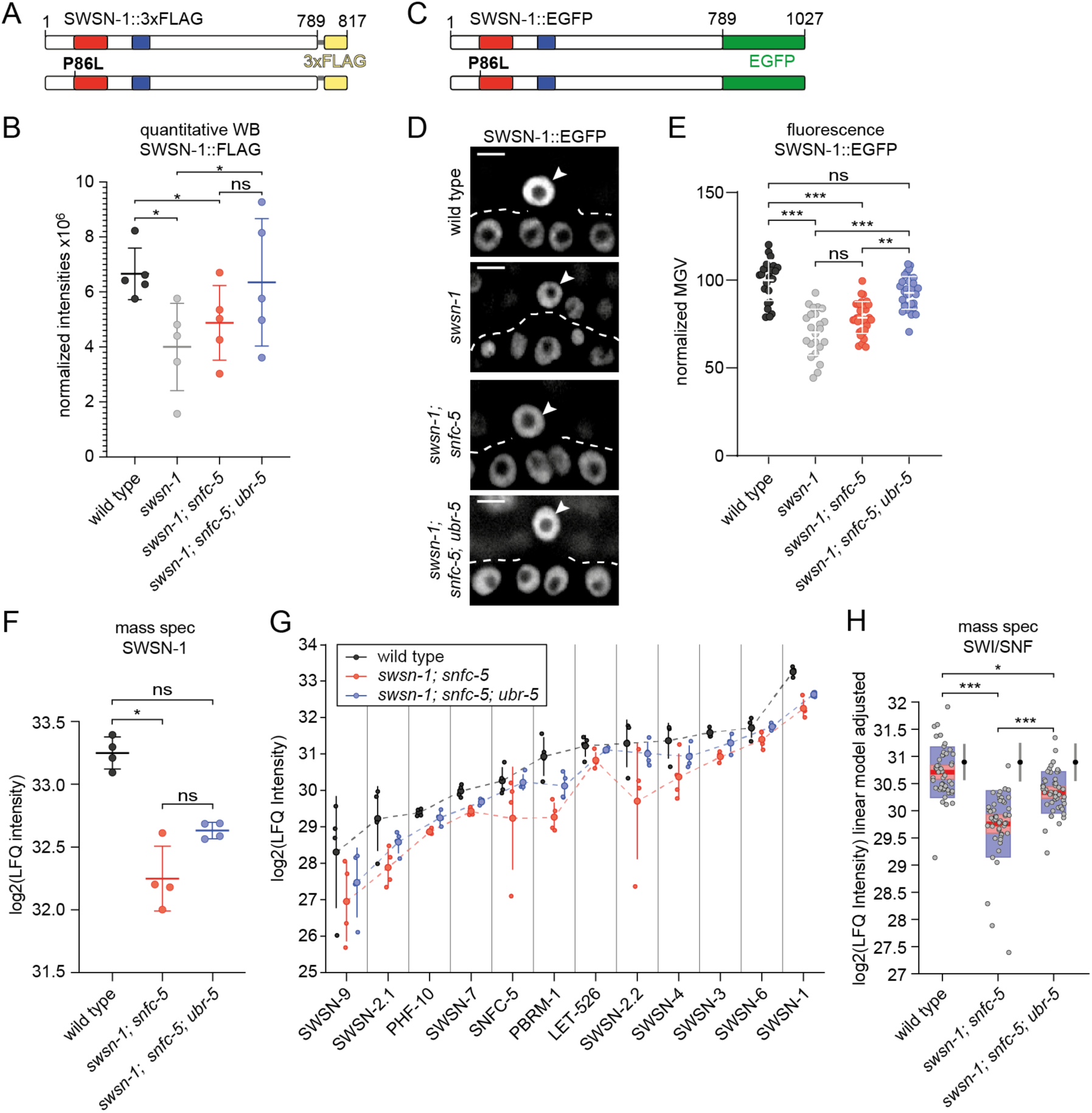
UBR-5 regulates SWI/SNF protein levels. **A)** In scale schematic representation of the wild-type and P86L SWSN-1::FLAG proteins. The SWIRM protein domain is shown in red, the SANT protein domain in blue, the glycine linker in grey and the 3xFLAG-tag in yellow. **B)** Western blot quantification of SWSN-1::FLAG protein levels in synchronized L1-staged wild-type (*swsn-1::3xflag*), *swsn-1* single mutant (*swsn-1^ts^::FLAG)*, *swsn-1; snfc-5* double mutant *(swsn-1^ts^::3xflag; snfc-5)* and *swsn-1; snfc-5; ubr-5* triple mutant *(swsn-1^ts^::3xflag; snfc-5; ubr-5)* animals (n=5). The y-axis represents raw integrated densities of SWSN-1::FLAG signals x10^6^ normalized to total protein signal. RM one-way ANOVA, * = Tukey’s multiple comparisons test adjusted p-value < 0.05, ns = not significant. **C)** In scale schematic representation of the wild-type and P86L SWSN-1::EGFP proteins. The SWIRM protein domain is shown in red, the SANT protein domain in blue and the EGFP-tag in green. **D-E)** Quantification of SWSN-1::EGFP intensities in the AC of P6.p 4-cell-staged wild-type (*swsn-1::egfp*), *swsn-1* single mutant (*swsn-1^ts^::egfp)*, *swsn-1; snfc-5* double mutant *(swsn-1^ts^::egfp; snfc-5)* and *swsn-1; snfc-5; ubr-5* triple mutant *(swsn-1^ts^::egfp; snfc-5; ubr-5)* animals (D) Representative images of quantified animals, ACs are indicated by white arrowheads, scale bar = 5µm. (The corresponding DIC images can be found in Figure S3A.) (E) SWSN-1::EGFP mean grey values (MGV) of ACs of individually quantified animals, Kruskal-Wallis test (H_3_ = 60.09, p < 0.0001), Dunn’s multiple comparison test adjusted p-value ** = < 0.001, *** = < 0.0001, ns = not significant. **F)** SWSN-1 protein levels in L1-staged wild-type, *swsn-1; snfc-5* double mutant and *swsn-1; snfc-5 ubr-5* triple mutant animals determined by label-free proteomics mass spec quantification (n=4). The y-axis represents log2 label-free quantification (LFQ) intensities. Kruskal-Wallis test (H_2_ = 8.769, p < 0.0012), Dunn’s multiple comparison test adjusted p-value * = < 0.01, ns = not significant. **G)** Protein levels of SWI/SNF subunits in synchronized L1-staged wild-type (grey), *swsn-1; snfc-5* double mutant (red) and *swsn-1; snfc-5 ubr-5* triple mutant (blue) animals determined by mass spec using label-free quantification (n=4). Small dots show protein levels of individual replicates, large dots indicate the mean protein levels and vertical lines the 95% confidence intervals. The dashed lines connect the mean protein levels of the different subunits for better visualisation of the overall trend. The y- axis represents log2 label-free quantification (LFQ) intensities. **H)** Box plot representation of the SWI/SNF complex protein levels adjusted for subunit type by multiple linear regression (see Figure S3B-C) data of the twelve SWI/SNF subunits from G combined. Red horizontal lines represent the median, light red boxes the 95% confidence intervals of the median and the blue boxes represent the standard deviation. The y-axis represents log2(LFQ intensities) of each genotype adjusted by subunit based on the linear model. F-Test, Bonferroni multiple comparison test adjusted p-value * < 0.01, *** < 0.0001. The intervals to the right of the boxplots show the mean (black dots) and standard deviation (blue lines) of 100 randomly chosen sets of 12 proteins. Alleles used: *swsn-1(ku355), swsn-1(syb2756[swsn-1::3xflag]), swsn-1(mj660; syb2756[swsn-1::3xflag]), swsn-1(st12187[swsn-1::egfp]), swsn-1(mj661; st12187[swsn-1::egfp]), snfc-5(mj633), ubr-5(mj638)*.

As an alternative approach to assess SWSN-1 protein levels at the single cell level in the different mutant animals, we measured SWSN-1::EGFP levels in the in the AC of mid-late L3-staged animals that had been exposed to the restrictive temperature (25°C) from the L2-L3 molt/early L3 stage until the P6.p 4-cell stage by confocal fluorescence microscopy. For this purpose, we obtained an available SWSN-1::EGFP strain, in which we introduced the P86L temperature-sensitive mutation (Figure 3C) and crossed it to *snfc-5* and *ubr-5* mutants to generate double and triple mutants. Consistent with the Western blot quantifications, we found that SWSN-1 protein levels in the AC were significantly reduced in *swsn-1* single and *swsn-1; snfc-5* double mutants compared to wild-type animals (Figure 3D-E and S3A). In this cell specific context, we observed a statistically significant increase of SWSN-1 levels in the in *swsn-1; snfc-5; ubr-5* triple mutants when compared to *swsn-1* single and *swsn-1; snfc-5* double mutants (Figure 3D-E and S3A), indicating that UBR-5 regulates SWSN-1 levels in the AC.

To gain better insight into the role of UBR-5 in regulating the abundance of SWI/SNF subunits, we employed a mass spectrometry approach to look at alterations in protein levels in wild-type animals, *swsn-1; snfc-5* double mutants and *swsn-1; snfc-5; ubr-5* triple mutants. Label-free quantification (LFQ) of whole proteome samples from synchronized L1-staged animals revealed a significant decrease in SWSN-1 protein levels in *swsn-1; snfc-5* double mutants compared to wild type animals (Figure 3F). Consistently with Western blotting results, SWSN-1 levels were partially restored to wild-type levels in *swsn-1; snfc-5; ubr-5* triple mutants (Figure 3F), a trend that was consistent in all detected SWI/SNF subunits (Figure 3G). Notably, every subunit of the complex showed this same pattern; the highest levels were in the wild type animals, reduced levels in the *swsn-1; snfc-*5 double mutants, and partially recovered levels in the *swsn-1; snfc-5; ubr-5* triple mutants, however, the mean protein abundance of each subunit was different. Therefore, we performed a multiple linear regression analysis to determine the effect of the mutation on the entire complex by regressing out the effect of the different mean expression levels. This analysis revealed that the SWI/SNF complex, as a whole, is significantly depleted in the *swsn-1; snfc-*5 double mutants when compared to wild-type animals and that this depletion is partially rescued in *swsn-1; snfc-5; ubr-5* triple mutants (Figure 3H and S3B-C). The same analysis with 100 randomly chosen sets of twelve proteins was not significant (depicted by the intervals to the right of the boxplots in Figure 3H). Figure S3D-E shows an example of a regression analysis for one set of twelve randomly selected proteins (ARX-6, EXOS-1, CYN-1, NBET-1, EMB-4, ZK1236.5, VPS-29, TFTC-3, MDT-9, TBA-1, HMT-1 and Y39G8B.1). All together these data indicate that the loss of UBR-5 results in increased protein levels of all SWI/SNF subunits in the *swsn-1; snfc-5* mutant model. Furthermore, these results suggest that UBR-5 likely ubiquitinates the SWI/SNF complex to tag it for proteasomal degradation and that the loss of UBR-5 likely prevents some of the developmental defects exhibited in SWI/SNF mutants by increasing SWI/SNF complex abundance. Our findings that levels of all SWI/SNF subunits of *swsn-1; snfc-5* double mutants are reduced compared to wild-type animals are consistent with previously published mouse data (Narayanan et al., 2015).

To determine how UBR-5 and HECD-1 regulate protein levels in animals, we similarly obtained mass spec proteomics data of wild-type animals, *ubr-5* single and *hecd-1* single mutants. When comparing protein levels of SWI/SNF subunits, we found that SWSN-7 and PHF-10 are significantly upregulated (FDR < 0.05) by 11 and 23% respectively in *ubr-5* single mutants compared to wild-type animals (Table S1) and that SWSN-3 is significantly upregulated (FDR < 0.05) by 10% in *hecd-1* single mutants compared to wild-type animals (Table S2). This data suggests that the steady state levels of specific individual SWI/SNF subunits could also be regulated by UBR-5 and HECD-1 ubiquitination, but that the loss of UBR-5 or HECD-1 does not broadly increase the abundance of all SWI/SNF subunits in wild-type animals.

It remains possible that in addition to regulating SWI/SNF protein levels, UBR-5 also ubiquitinates other proteins and that restoring the levels of these proteins also helps SWI/SNF mutants develop to adulthood. Our proteomics analysis identified twelve significantly upregulated proteins (fold-change > = 1.5: FDR <= 0.05) (SKP-1, F08F3.4, FBXA-156, ZK228.4, T19H12.2, F36A2.3, GST-41, PCN-1, HAT-1, HRG-2, HSP-16.1 and HSP-16.49) in *swsn-1; snfc-5; ubr-5* triple mutants when compared to *swsn-1; snfc-5* double mutants (Figure S3F). The collective upregulation of those twelve proteins in *swsn-1; snfc-5* mutants might contribute to the suppression of observed developmental defects. Furthermore, using the proteomics data of *ubr-5* and *hecd-1* single mutants, we could identify 21 proteins significantly upregulated in both mutant conditions (Figure S3G), suggesting that the regulation of those proteins might be a redundant function of the two ubiquitin ligases.

### Mutations that suppress the embryonic lethality of *swsn-1* mutants restore wild-type expression of 335 SWI/SNF regulated genes

As chromatin remodellers, the SWI/SNF complexes are thought to control the expression of various genes by enabling or preventing DNA access for the transcription machinery. This involves positioning of nucleosomes to expose promoter and enhancer sequences, thereby enabling the binding of transcription factors and RNA polymerase II (RNA pol II). Similarly, SWI/SNF complexes can facilitate the binding of repressors and disable the access to transcription start sites (TSS) (Saha et al., 2006; Wilson and Roberts, 2011). Previous studies analysing the transcriptomes of young adult-staged *swsn-1* mutants, reported that between 7.5% (Riedel et al., 2013) and approximately one third (Mathies et al., 2020) of *C. elegans* genes are regulated by the SWI/SNF complex. To identify changes in gene expression that might lead to the developmental defects of *swsn-1* mutants, we conducted three independent RNA-sequencing experiments (sets 1 – 3) using synchronized L1-staged animals (same conditions as for the quantitative Western blotting and proteomics experiments). We analyzed gene expression profiles of wild-type animals, *ubr-5* mutants, *swsn-1* mutants, and various combinations of double and triple mutant animals to investigate how these different mutations affect SWI/SNF regulated gene expression. We found that approximately 8% of *C. elegans* protein coding genes (1803) were differentially expressed in *swsn-1* single mutants compared to wild-type animals (Figure 4A). The expression changes of most of these genes were restored to wild-type levels in the *swsn-1; snfc-5* double, *swsn-1; snfc-5; ubr-5* triple and *swsn-1; snfc-5; hecd-1* triple suppressor mutants but not restored in *swsn-1; ubr-5* double mutants (Figure 4A). The gene expression profiles of *swsn-1; ubr-5* double mutants were similar to those of *swsn-1* single mutants (Figure S4A) which is consistent with our previous findings that both of these mutant backgrounds exhibit embryonic lethality under the stringent heat-shock conditions (Figure 1A-B). These data are consistent with our hypothesis that the *snfc-5* A258V mutation generally suppresses the defects caused by the *swsn-1* mutation and that the *ubr-5* mutation alone is not sufficient to suppress defects when animals are exposed to the restrictive temperature at early larval stages. *ubr-5* single mutants had only 62 differentially expressed genes compared to wild-type animals (Figure 4A).

**Figure 4:**
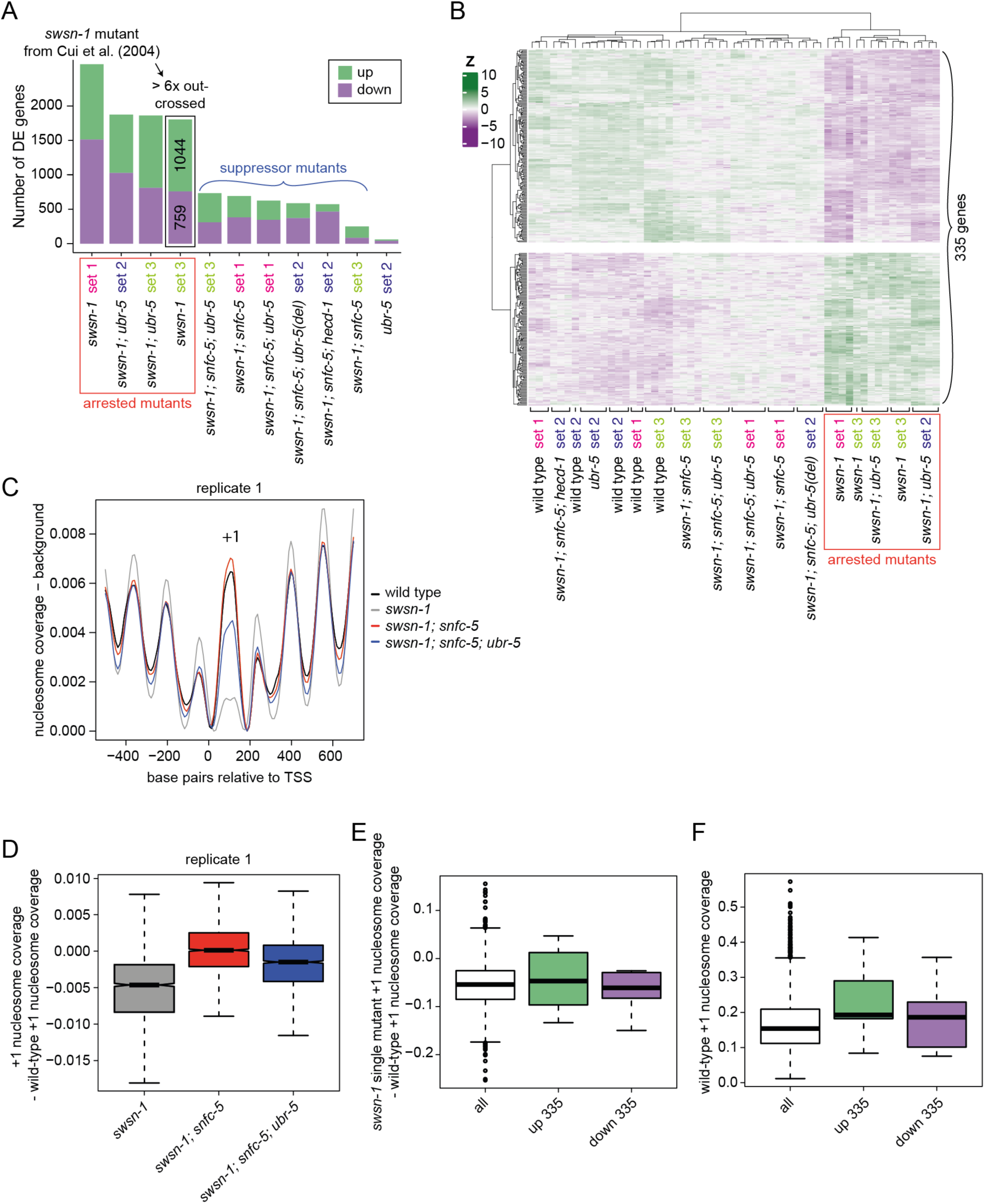
Mutations that suppress the embryonic lethality of *swsn-1* mutants restore wild-type expression of 335 SWI/SNF regulated genes. **A-B)** Differential gene expression analysis of three independent RNA-sequencing (RNA-seq) experiments (sets 1 -3) using synchronized L1-staged wild-type animals and *swsn-1* single, *swsn-1; snfc-5* double, *swsn-1; snfc-5; ubr-5* triple (Q150* and deletion alleles), *swsn-1; snfc-5; hecd-1* triple, *swsn-1; ubr-5* double and *ubr-5* single mutants (n >= 4). (A) Bar graphs of differentially up- (green) or downregulated (purple) genes of the different mutants compared to wild-type animals of their respective RNA-seq set. (B) Z-score heatmap of 335 genes consistently differentially expressed (DE) only in *swsn-1* single and *swsn-1; ubr-5* double mutants. **C-F)** Nucleosome coverage analysis of MNase-sequencing (MNase-seq) data using synchronized L1-staged wild-type animals and *swsn-1* single, *swsn-1; snfc-5* double and *swsn-1; snfc-5; ubr-5* triple mutants. (C) Nucleosome traces around the TSS of ubiquitous genes. (D) Box plots of locus-by-locus +1 nucleosome coverage of ubiquitous genes in the mutants relative to wild-type coverage. Bold horizontal lines represent the median, boxes represent interquartile range and whiskers extend to the greatest point <=1.5 times the interquartile range. (E) Box plots of *swsn-1* single mutant +1 nucleosome coverage minus wild-type +1 nucleosome coverage of all ubiquitous genes (white) and ubiquitous up- (green) and downregulated (purple) genes from the 335 genes from B. Bold horizontal lines represent the median, boxes represent interquartile range and whiskers extend to the greatest point <=1.5 times the interquartile range. Individual data points represent outliers. (F) Box plots of the +1 nucleosome coverage in wild-type animals of all ubiquitous genes (white) and ubiquitous up- (green) and downregulated (purple) genes from the 335 genes from B. Bold horizontal lines represent the median, boxes represent interquartile range and whiskers extend to the greatest point <=1.5 times the interquartile range. Individual data points represent outliers. Alleles used: *swsn-1(ku355), snfc-5(mj633), ubr-5(mj638), ubr-5(ok1108)* (used in *swsn-1; snfc-5; ubr-5* triple mutant from RNA-seq set 2, indicated by ‘(del)’)*, hecd-1(ok1437)*.

We hypothesized that the suppressor mutants we identified restore developmental defects of *swsn-1* mutants by restoring the expression of developmentally critical genes. However, identifying the precise SWI/SNF regulated genes that promote animal development is complicated by the asynchronous developmental rate of SWI/SNF mutant animals (see developmental progression estimation for the sequenced animals using an available RNA-seq time-course data set in Figure S4B), which can be slower than wild-type animals. To not confound developmentally important genes with genes that are differentially expressed due to developmental staging differences, we compared differentially expressed genes between mutant backgrounds that exhibited early developmental arrest (*swsn-1* single mutants and *swsn-1; ubr-5* double mutants, see Figure S1G) and genetic backgrounds that did not exhibit early developmental arrest (wild-type animals, *swsn-1; snfc-5* double mutants, *swsn-1; snfc-5; ubr-5* triple mutants, *swsn-1; snfc-5; hecd-1* triple mutants, and *ubr-5* single mutants) at the developmentally synchronized L1 larval stage (i.e. all genetic backgrounds were synchronized as L1 stage animals). Furthermore, we performed these RNA-seq experiments in multiple different “sets” (each with four or five biological replicates) and focused only on genes exhibiting consistent expression changes in all experiments. From this filtered analysis, we identified 335 genes (out of the 1,083 genes differentially expressed in *swsn-1* single mutants when compared to wild-type animals) that were consistently differentially expressed in the strongly developmentally arrested *swsn-1* single and *swsn-1; ubr-5* double mutants (Figures 1A-B and S1D-H) but displayed near wild-type gene expression in all mutant backgrounds that did not exhibit the early larval arrest (Figure 4B). We conclude that the dysregulation of this subset of genes is likely at least partially responsible for driving the early developmental arrest that occurs in *swsn-1* single mutants.

To evaluate the potentially direct targets of altered SWI/SNF function on nucleosome positioning at gene promoters, we performed MNase-sequencing of synchronized L1-staged (same conditions as for RNA-seq) wild-type animals, *swsn-1* single, *swsn-1; snfc-5* double and *swsn-1; snfc-5; ubr-5* triple mutants to obtain genome-wide nucleosome coverage profiles. MNase digests in nucleosome-free regions resulting in a higher signal corresponding to regions protected by nucleosomes, which we refer to as higher nucleosome coverage. This can result from increased nucleosome density (i.e. increased probability of a nucleosome being present at all) or a more precisely positioned nucleosome (i.e. the nucleosome is less “fuzzy”) (Cui and Zhao, 2012). Since only promoters of ubiquitous and germline-specific genes have clear and well-positioned nucleosomes (Serizay et al., 2020) and we sequenced *C. elegans* L1 larvae that do not have extensive germline tissue (Sulston and Horvitz, 1977), the MNase-seq analysis was restricted to ubiquitous genes. The clearest difference we observed between wild-type animals and *swsn-1* single mutants was a strongly reduced +1 nucleosome coverage (the first nucleosome downstream of the promoter). The +1 nucleosome coverage in both the *swsn-1; snfc-5* double and *swsn-1; snfc-5; ubr-5* triple suppressor mutants was higher than in *swsn-1* single mutants (Figure 4C and S4C). These findings indicate that SWI/SNF regulates the nucleosomes at the +1 position of ubiquitously expressed genes and that our suppressor mutations can partially restore the defects observed in *swsn-1* single mutants. Notably, the reduction of the +1 nucleosome coverage in *swsn-1* mutants relative to wild-type nucleosome coverage can also be seen comparing across all ubiquitous promoters separately, indicating a consistent difference that affects most ubiquitous promoters (Figure 4D and S4D). The reduced coverage could reflect less well-positioned +1 nucleosomes or reduced nucleosome density at the +1 region. These data suggest that mutation of *swsn-1* affects the chromatin remodelling function of the SWI/SNF complex, which might in turn explain the observed changes in gene expression.

We next asked if our set of 335 developmentally regulated SWI/SNF genes (defined in Figure 4B) has a distinct nucleosome positioning profile. To do so, we first asked whether genes with altered expression showed larger changes in nucleosome coverage than genes that did not show changes in expression. We compared the +1 nucleosome coverage of all ubiquitous genes to ubiquitously up- (green) and downregulated (purple) developmentally regulated SWI/SNF genes, which are subsets of the 335 genes. This analysis revealed that the promoters of ubiquitous genes altered in *swsn-1* mutants had a similar reduction in +1 nucleosome coverage relative to wild-type animals (Figure 4E). These results suggest that the coverage of the +1 nucleosome at gene protomers changes for many genes in *swsn-1* single mutants, but this does not always result in changes in their expression.

Interestingly, we found that the subsets of ubiquitously up- (green) and downregulated (purple) SWI/SNF-dependent genes have a higher nucleosome coverage in wild-type animals when compared to all ubiquitous genes in wild-type animals (Figure 4F). Figure 4E indicates that *swsn-1* mutants also have a higher nucleosome coverage in this subset of genes. These findings show that the 335 developmentally important SWI/SNF genes tend to have more well positioned +1 nucleosomes or a higher nucleosome density than (ubiquitous) genes on average. This could suggest that developmentally regulated SWI/SNF genes depend on a well-positioned +1 nucleosome or on a high nucleosome density for their regulation, and that other genes might be more robust to changes in nucleosome coverage.

## Discussion

Here we identified that the combination of a specific missense mutation in *snfc-5* and the loss of either of two E3 ubiquitin ligases (UBR-5 or HECD-1) can suppress some of the developmental defects caused by a missense mutation in the core SWI/SNF subunit *swsn-1.* UBR-5 and HECD-1 are novel genetic interactors of the SWI/SNF complex, and our studies revealed a previously unknown functional redundancy between these two ubiquitin ligases in regulating animal development and fertility. In addition, we established that *swsn-1; snfc-5* double mutants have reduced SWI/SNF protein levels, and that the loss of UBR-5 can partially restore these levels. Together, these results are consistent with a model in which UBR-5 and HECD-1 promote the degradation of the SWI/SNF complex and that the loss of UBR-5 or HECD-1 can prevent developmental defects from manifesting by increasing SWI/SNF levels (Figure 5). Missense mutations in SWI/SNF complex subunits cause a multitude of developmental disorders in humans. Heterozygous mutations identified in developmental disorders are generally dominant, implying that dosage-sensitive processes underlie the roles of SWI/SNF complexes in development (Kadoch and Crabtree, 2015). Our findings suggest that the inhibition of UBR5 and HECTD1 could be a viable therapeutic strategy to treat developmental disorders caused by dosage sensitive missense mutations in SWI/SNF complex subunits.

**Figure 5:**
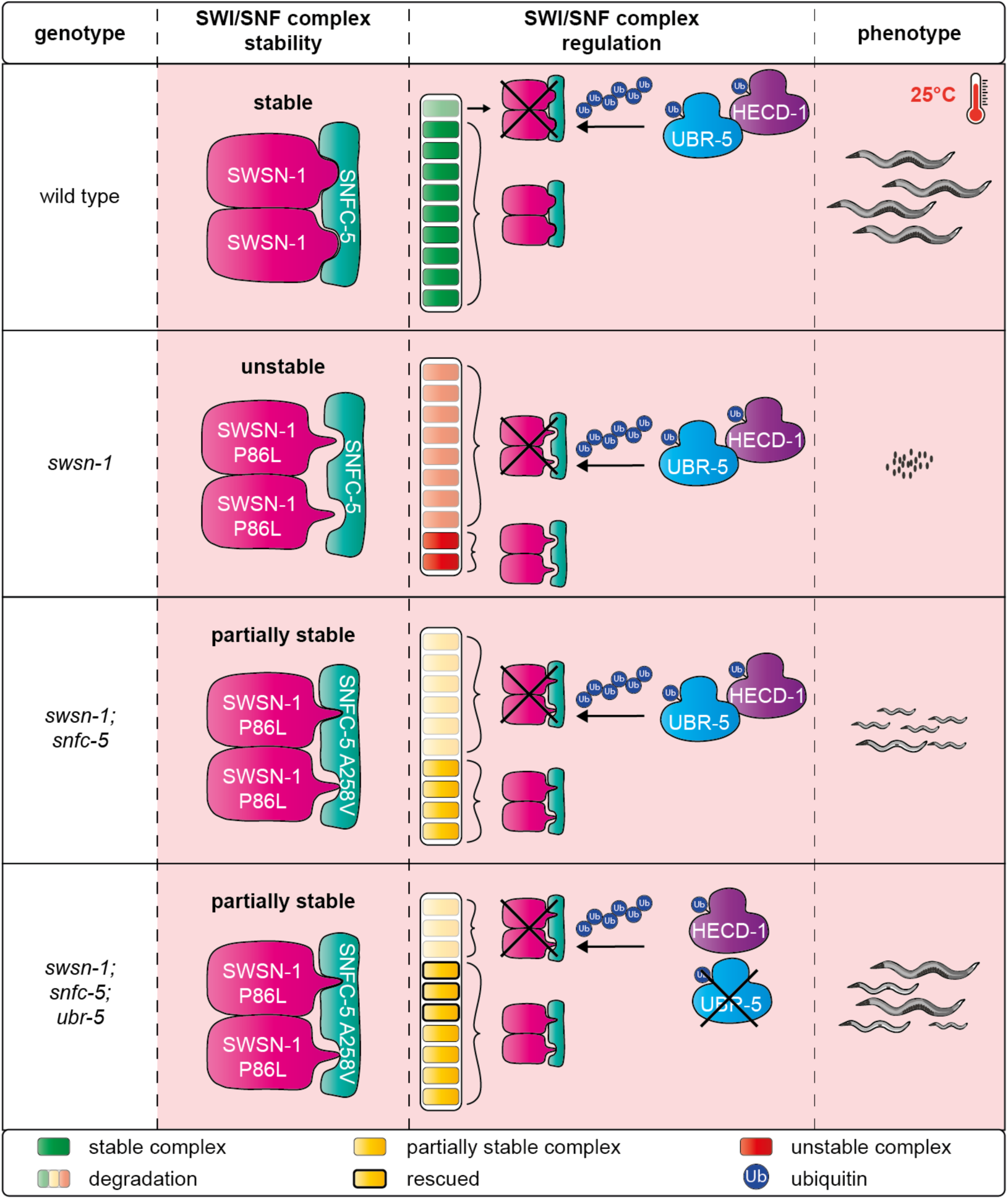
Model of *swsn-1* mutation suppression by *snfc-5* and *ubr-5* mutation. In wild-type animals, a stable SWI/SNF complex can assemble at 25°C (indicated by a pink box and a thermometer). UBR-5 and HECD-1 ubiquitinate SWI/SNF complexes for proteasomal degradation to maintain their steady state levels. The *swsn-1* mutation prevents the assembly of the SWI/SNF complex or leads to the assembly of an unstable complex at 25°C due to structural changes that impair the interaction of SWSN-1 and SNFC-5. Mutant SWI/SNF complexes are more frequently degraded by the proteasome, which is also mediated by UBR-5 and HECD-1 ubiquitination. Additional mutation of *snfc-5* enables the assembly of a partially stable mutant SWI/SNF complex that gets overall less frequently degraded by the proteasome. Loss of UBR-5 leads to an increase of SWI/SNF complex protein levels in *swsn-1; snfc-5* mutants by preventing some of the turnover of the complex by the proteasome.

### SWI/SNF complex stability

The human SMARCC1/2 dimer is important in early SWI/SNF complex assembly where it serves as a scaffold to which other subunits bind (He et al., 2020; Mashtalir et al., 2018). Studies of SWI/SNF complex assembly found that loss of SMARCC1/2 in mice leads to the dissociation and degradation of the entire complex, presumably because unassembled subunits are less stable (Narayanan et al., 2015). In the SWI/SNF mutant *C. elegans* model we used here, a P86L mutation in the SWIRM domain of the *SMARCC1/2* homolog *swsn-1* resulted in embryonic lethality at 25°C. We rescued the embryonic lethality by introducing a A258V mutation in the RPT2 domain of *snfc-5*, homolog of *SMARCB1*. The SWIRM domains of human SMARCC1/2 and yeast SWI3 dimers directly bind the RPT1 and RPT2 domains of SMARCB1 and SNF5 respectively (Han et al., 2020; He et al., 2020). Based on these data, we propose a model in which the P86L SWSN-1 mutation results in a structural change which prevents stable binding to SNFC-5 and complex assembly. This is supported by the fact that the allele is temperature-sensitive, as we expect destabilization to be exacerbated by increased temperature. The A258V SNFC-5 mutation in turn introduces another structural change that allows SNFC-5 to bind mutant SWSN-1 more stably (Figure 5). The resulting *swsn-1; snfc-5* mutant complex is partially stable, but not as stable as the wild-type assembly, since we find that the levels of all SWI/SNF subunits in *swsn-1; snfc-5* double mutants are reduced compared to wild-type animals (Figure 3G-H). Consistent with this, it has previously been proposed that loss of one SWI/SNF subunit could alter the abundance of the other subunits (Euskirchen et al., 2011).

We also showed that the levels of all SWI/SNF subunits are partially restored to wild-type levels in *swsn-1; snfc-5; ubr-5* triple mutants (Figure 3G-H). Moreover, even though the loss of either UBR-5 or HECD-1 did not substantially affect SWI/SNF levels in wild-type animals, we found that some individual SWI/SNF subunits were increased by approximately 10-20% in *ubr-5* and *hecd-1* single mutants (Tables S1 and S2). This suggests that UBR-5 and HECD-1 can also affect the steady state levels of wild-type SWI/SNF subunits. Overall, our data are consistent with a model in which UBR-5 (and presumably also HECD-1) directly ubiquitinates either SWSN-1 or another subunit of the SWI/SNF complex to regulate SWI/SNF protein levels. Specifically, our data suggest that *swsn-1* single mutants form an unstable SWI/SNF complex that consequently gets degraded by the proteasome resulting in very low SWI/SNF protein levels and embryonic lethality. By contrast, *swsn-1; snfc-5* double mutants form a partially stable complex that is less prone to degradation resulting in increased SWI/SNF complex abundance, when compared to *swsn-1* single mutants, and the prevention of embryonic lethality. Lastly, the loss of UBR-5 or HECD-1 further increase SWI/SNF complex abundance in *swsn-1; snfc-5* double mutants by preventing some of the turnover of the complex by the proteasome. This ultimately promotes SWI/SNF function and enables *swsn-1; snfc-5; ubr-5* triple mutants to develop to adulthood (Figure 5).

### Synergistic functions of UBR-5 and HECD-1

We identified two paralogous HECT-type E3 ubiquitin ligases as suppressors of SWI/SNF mutation in our screen. Despite the finding that the loss of either of the two enzymes alone appears to promote animal health in *swsn-1; snfc-5* mutants, we found that the combined loss of these two proteins has deleterious effects on animal health. To our knowledge, this is the first observation of a synergistic genetic interaction between these two ubiquitin ligases. These findings suggest that UBR-5 and HECD-1 likely function at least partially redundantly. Our data indicates that the SWI/SNF complex is one likely such common target of UBR-5 and HECD-1, however our proteomics profiling of *ubr-5* and *hecd-1* single mutants only observed small increases of different individual SWI/SNF subunits. We suspect that due to the potentially redundant functions of UBR-5 and HECD-1, there would be substantially larger changes in SWI/SNF subunit abundance and other protein abundances in *ubr-5; hecd-1* double mutants which cannot properly regulate protein levels via either ubiquitin ligase. These dysregulated protein levels might in turn drive the strong synthetic phenotypes observed in *ubr-5; hecd-1* double mutants.

### UBR-5 function at later developmental time points

Despite our finding that *swsn-1; snfc-5; ubr-5* triple mutants could develop to adulthood, making them better developmental suppressors than the *swsn-1; snfc-5* double mutants, we did not observe an improved rescue of gene expression or of nucleosome coverage in the triple mutants when compared to double mutants. We suspect that the loss of UBR-5 on SWI/SNF-dependent gene expression likely becomes more apparent at later developmental stages. This is supported by our observations that *ubr-5* alone, in the absence of any *snfc-5* mutation, was capable of partially suppressing *swsn-1* single mutants if they survived to a later (post L1) larval stage (Figure 1D-E and S1K). Future studies looking at different developmental time points might shed light on how and if UBR-5 affects the gene expression of SWI/SNF target during development.

### Evolution of SWI/SNF and complex stability

Interestingly, many animals including humans, mice, chicken, zebrafish, and fruit flies all already have a valine at the equivalent position as the A258V substitution in the RPT2 domain of SNFC-5 that we identified as a suppressor in the first mutagenesis screen (Figure S1C). Moreover, one of the mutants isolated in the second mutagenesis screen carries the additional V129I substitution mutation in the SWIRM domain of *swsn-1* and the human, mouse and fruit fly SWSN-1 homologs also have an isoleucine at the equivalent position (Figure S1A). These findings suggest the valine and isoleucine at these specific positions could make the SWI/SNF complex more robust and these exact substitutions might have played a role in the evolution of SWI/SNF function. For example, these specific substitutions might help animals better adapt to higher temperatures such as those found in warm blooded animals or even the poikilotherm model animals zebrafish and fruit flies which develop at approximately 28°C (Reed and Jennings, 2011) and 25°C (Hamada et al., 2008) respectively.

## EXPERIMENTAL MODEL DETAILS

### *C. elegans* strain maintenance

*C. elegans* strains were grown and maintained on nematode growth medium (NGM) agar plates seeded with HB101 bacteria (Caenorhabditis Genetics Center, University of Minnesota, Twin Cities, MN, USA) as food source (Brenner, 1974). Strains containing the temperature-sensitive *swsn-1* mutation were routinely kept at 15°C unless otherwise indicated and other strains were routinely kept at 20°C or also at 15°C to match growth conditions of temperature-sensitive strains. The *C. elegans* strains that were used in this study are derived from the Bristol N2 strain and are listed in the key resource table.

## METHOD DETAILS

### EMS mutagenesis screening

Ethyl methosulfate (EMS) mutagenesis was performed as described previously by Brenner (Brenner, 1974). Larval 4 (L4)-staged *C. elegans* were washed off plates with M9 buffer, collected in 15ml falcon tubes and washed three times with M9 buffer. Animals were then incubated with a final concentration of 0.05M ethyl methosulfate (Sigma Aldrich) in 4ml M9 buffer on a rotating wheel for four hours at room temperature. After four washes with M9 buffer animals were seeded onto several agar plates and left to recover at 20°C. Once the F1 offspring of mutagenized animals were gravid adults, F1 animals were bleached to obtain F2 generation animals. L4 and young adult staged F2 animals were either maintained at 20°C (first screen) or shifted to 25°C for approximately 16 hours (second screen) and the then gravid adult F2 animals were bleached to obtain F3 generation embryos. The F3 embryos were grown at 25°C for three to five days and screened for mutants that developed to adulthood. Individually picked F3 mutant animals with the desired phenotype were recovered at 15°C.

### Bleaching and synchronisation of *C. elegans*

Gravid adult *C. elegans* were washed off NGM plates in M9 buffer (0.6% Na_2_HPO_4_, 0.3% KH_2_PO_4_, 0.5% NaCl and 1mM MgSO_4_) and collected in 15ml falcons or 1.5ml tubes. Animals in 15ml tubes were pelleted by centrifugation at 2000rpm for 1 minute and M9 buffer was aspirated to leave 2ml, animals in 1.5ml tubes were pelleted by centrifugation at 3000rpm for 30 seconds and M9 buffer was aspirated to leave 0.5ml. An equal volume of 2x bleaching solution (1M NaOH and 1.5% NaClO - free chlorine) was added and animals were vortexed vigorously for 4-6 minutes to destroy adults and recover embryos. Embryos were washed at least twice in M9 buffer and either directly seeded onto new NGM plates or left to hatch in 5ml M9 rotating on a wheel at 20°C for 24 hours to obtain a synchronized population of L1s.

### Collection of synchronized L1-staged *C. elegans*

L1s that hatched after bleaching were counted three times in 10µl drops of M9 buffer to estimate the number of animals present in the 15ml tubes. After 24 hours rotation at 20°C, tubes were spun down at 4000g for 1 minute, M9 buffer aspirated and a certain number of L1s seeded on 9cm NGM agar plates, depending on the assay. At least 30,000 L1s per sample were seeded for Western blotting, 100,000 to 200,000 L1s were seeded for proteomics, 10,000 to 30,000 L1s were seeded for RNA-sequencing and 55,000 to 60,000 L1s were seeded for MNase-sequencing. Subsequently, L1s were grown at 25°C for six hours, a period after which the animals are still at the larval 1 stage of development. L1s were collected into 15ml tubes in M9 buffer with P1000 tips coated in 0.05% TWEEN-20 (Sigma Aldrich) in M9 buffer. Animals were washed three times in 15ml M9 buffer by pelleting animals with 1-minute centrifugations at 4000g and aspiration of buffer.

After the last wash, L1s to be used for Western blotting, proteomics and RNA-sequencing were transferred into 1.5ml tubes with tips coated in 0.05% TWEEN-20 (Sigma Aldrich) in M9 buffer, spun 1 minute at 8000g and remaining M9 buffer was aspirated carefully to leave little buffer on the pellets. For protein extractions, 100µl pellets of L1s in M9 were snap frozen with liquid nitrogen and stored at −70°C. For RNA extractions, 500µl Trizol reagent (Thermo Fisher Scientific) was added to the pellets and samples stored at −70°C.

For MNase-sequencing, small frozen “worm balls” were generated from the L1 pellets containing little M9 buffer. This was done by dripping small amounts of animal/M9 buffer mix into a cooled ceramic bowl placed on top of dry ice and filled with liquid nitrogen using a glass Pasteur pipette. “Worm balls” were carefully collected into 1.5 ml tubes with a cooled metal spoon and stored at −70°C.

### CRISPR-Cas9 gene editing

CRISPR-Cas9 gene editing was performed essentially as described previously (Paix et al., 2015). For the injection mixes 0.5µl KCl (0.5M), 0.74µl Hepes pH 7.5 (100mM), 2.5μl tracrRNA (4 μg/μl, Dharmacon), 0.4μl target gene gRNA (4μg/μl, Dharmacon), 0.4μl homologous recombination repair template (1 μg/μl, IDT) and 50ng Pmyo-3::mCherry::unc-54 co-injection plasmid (pCFJ104) (Frøkjær-Jensen et al., 2008) were mixed. Then 0.75µl Cas9 (2.5µ/µl, Dharmacon) and DEPC water up to a volume of 10μl were added, mixed, and incubated at 37°C for 15 min. The mix was spun down at max. speed in a tabletop centrifuge, 7.5µl of the mix was transferred into a new tube and micro-injected into the germline young adult staged animals. For the injections, animals were transferred into a drop of halocarbon oil 700 (Sigma Aldrich) on a cover slip with a 2% agarose pad. The animals were straightened in the oil drop using an eyelash pick and positioned so gonad arms could be injected with Femptotip injection capillaries (Eppendorf). An Olympus IX71 microscope equipped with a micromanipulator, FemtoJet injection rig and InjectMan joystick (Eppendorf) was used for the injections. After the injection, M9 buffer was added to the animals to remove the oil and the animals were transferred to individual plates and recovered at 20°C overnight. The offspring of injected animals was then screened for animals expressing the red co-injection plasmid in body wall muscles at a fluorescence microscope. From positive plates, approximately 100 F1 animals were singled onto new plates and genotyped for the introduced allele after they produced F2 offspring. F2 offspring of F1 animals carrying the heterozygous desired allele were singled again and genotyped to obtain a homozygous mutant. The gRNAs and repair templates that were used in this study are listed in the key resource table.

### Temperature shift assays

To assess temperature-sensitive developmental defects of different *swsn-1* mutants and wild-type animals, animals were synchronized by bleaching as described in section ‘Bleaching and synchronisation of *C. elegans*’. To assess developmental differences between *swsn-1* single and *swsn-1; snfc-5* double mutants, recovered embryos were directly seeded onto NGM agar plates and grown at 25°C for 48 hours (Figure S1D-F). To assess developmental differences between *swsn-1* single and *swsn-1; ubr-5* double mutants, recovered embryos were directly seeded onto NGM agar plates and grown at 22.5°C for 48 hours and 25°C for 24 hours (Figure 1D-E and S1K). To assess developmental differences between *swsn-1; snfc-5* double mutants and *swsn-1; snfc-5; ubr-5* triple mutants, more stringent conditions were used. Recovered embryos were directly seeded onto NGM agar plates, grown at 15°C until the L4 or young adult stage and then shifted to 25°C for 16 hours. Gravid adults were bleached and recovered embryos were seeded onto new NGM agar plates and grown at 25°C for 72 hours (Figures 1A-B, S1H, 2B-C and S2).

Animals were classified in categories (L1 or > L1, < gravid adult or gravid adult and < L4 or L4 +, respectively) and manually counted (3 replicates of at least 100 animals per genotype were counted). Subsequently, animals were washed off plates, collected in 1.5ml tubes with M9 buffer, washed twice in 1ml M9, pelleted and M9 buffer aspirated. To image animals, 18µl of animals in M9 were deposited onto a glass slide coated with a 2% agarose pad and paralyzed by adding 2µl 100mM tetramisole. A coverslip was added and mounted with transparent nail polish. Representative differential interference contrast (DIC) microscopy images of animals were taken with a Leica DM6B microscope (upright microscope with LAS X imaging system) at 10x magnification and 30ms exposure time or 4x magnification and 10ms exposure time.

### Live offspring counts

Animals were grown at 20°C and twenty L4-staged animals were individualized onto 50mm NGM agar plates per genotype. On the following day, each egg-laying adult animal was transferred onto a fresh plate. This was repeated approximately every 24 hours until the production of embryos ceased. Number of hatched live offspring was counted manually 24-72 hours after removing the parental adult. Statistical analyses and plotting of data were conducted using GraphPad Prism (v. 9.0.0).

### Embryonic lethality and larval hatching quantification

Synchronized embryos (see section ‘Bleaching and synchronisation of *C. elegans*’) were directly seeded onto NGM agar plates, grown at 15°C until the L4 or young adult stage and then shifted to 25°C for 16 hours. Gravid adults were bleached, recovered embryos were washed 5 times in M9 buffer and counted three times in 10µl drops of M9 buffer to estimate the number of embryos. Approximately 200 embryos were seeded onto 50mm NGM agar plates, counted manually to determine the precise number of embryos seeded and grown at 25°C. After 24 hours, hatched larvae were manually counted to determine the percentage of larval hatching or embryonic lethality. Five replicates of approximately 200 animals were counted per genotype and numbers combined for Fisher’s exact test p-value calculations. Statistical analyses and plotting of data were conducted using GraphPad Prism (v. 9.0.0).

### Anchor cell (AC) assays

#### Assessment of AC invasion

Synchronized animals were grown at 15°C until the L2 molt-early L3 stages, and then shifted to 25°C until the P6.p4-cell stage. AC was defined under differential interference contrast (DIC). An intact barrier under the AC was used to assess invasion. Wild-type invasion was defined as a breach as wide as the basolateral surface of the AC (Sherwood and Sternberg, 2003). AC invasion was scored at the P6.p 4-cell stage, when 100% of wild-type animals exhibit a breach in the BM (Sherwood and Sternberg, 2003).

#### Live-cell imaging and image quantification

Animals were mounted into a drop of M9 buffer on a 5% Noble agar pad containing approximately 10mM sodium azide anesthetic and topped with a coverslip. Several experiments were scored using epifluorescence visualized on a Zeiss Axiocam MRM camera, also mounted on an upright Zeiss AxioImager A2 and a Plan-Apochromat 100×/1.4 (NA) Oil DIC objective. Microscopy images were obtained on a Hamamatsu Orca EM-CCD camera mounted on an upright Zeiss AxioImager A2 with a Borealis-modified CSU10 Yokagawa spinning disk scan head (Nobska Imaging) using 488nm Vortran lasers in a VersaLase merge and a Plan-Apochromat 100×/1.4 (NA) Oil DIC objective. MetaMorph software (Molecular Devices) was used for microscopy automation. Images were processed using Fiji/ImageJ (v.2.1.0/1.53c) (Schindelin et al., 2012). Expression levels of SWSN-1::EGFP were measured by quantifying the mean grey value of AC nuclei, defined as the somatic gonad cell near the primary vulva. Background subtraction was performed by rolling ball background subtraction (size = 50). Statistical analyses and plotting of data were conducted using GraphPad Prism (v. 9.0.0). Figure legends specify what statistical test and p-value cut-off was used to determine statistical significance.

### Preparation of genomic DNA

Starved *C. elegans* were washed of 90mm NGM plates (one plate relatively full of *C. elegans* per strain) in M9 buffer and collected in 15ml tubes. Animals were washed twice in M9 buffer, samples transferred into 1.5ml tubes, pelleted, most of the buffer was removed and pellets stored at −80°C. Animals were lysed with 1ml Cell Lysis Solution (Qiagen) and samples thawed during this process. Five µl Proteinase K (20mg/ml, Thermo Fisher Scientific) was added and incubated at 55°C for approximately 3 hours at 600rpm shaking, until only embryos were left. Lysates were cooled to room temperature and inverted periodically. Then, 5µl RNase A solution (Thermo Fisher Scientific) was added, samples incubated at 37°C shaking for 1 hour and cooled one ice for 3 minutes. Subsequently, 333µl Protein Precipitation Solution (Qiagen) was added and samples were vortexed vigorously for 20 seconds at high speed. Samples were centrifuged for 10 minutes at 2000rcf and 600µl supernatant transferred into new tubes that were prefilled with 500µl isopropanol and inverted 50 times to mix. After a 2-hour incubation at −20°C, samples were centrifuged at maximum speed for 5 minutes, the supernatant aspirated carefully, and DNA pellet washed with 750µl 75% ethanol by inverting the tubes several times. Samples were centrifuged again at 2000rcf for 3 minutes and supernatant carefully removed with aspirator. Pellets were air dried and resuspended in 35µl water. DNA samples were stored at 4°C.

### Whole genome sequencing

#### Preparation of genomic DNA libraries

Five µl of genomic DNA sample was run on an 1% agarose gel to ensure that DNA fragments were of approximately 10kb sizes. DNA concentrations and A260/A280 rations were determined using a Nanodrop spectrophotometer (Thermo Fisher Scientific) and concentrations were determined again by Qubit HS dsDNA fluorometric quantification (Thermo Fisher Scientific) according to the manufacturer’s instructions. Thirty µl of 100-500ng DNA per sample were prepared. Multiplexed DNA libraries were generated using the Nextera DNA Flex library prep kit (catalog number 20018704, Illumina) and Nextera™ DNA CD Indexes (catalog number 20018707, Illumina) according to the manufacturer’s instructions. Libraries were quantified again by Qubit (Thermo Fisher Scientific) and quality control was performed by TapeStation run using a D1000 ScreenTape (Agilent). Samples were sequenced on a HISeq 1500 machine (Illumina) in paired end mode with 50bp read length.

#### Data processing and analysis

Raw DNA sequencing reads were trimmed for low quality and adapters using Trimmomatic (Bolger et al., 2014) (Version 0.39, parameters: ILLUMINACLIP: TruSeq3-SE.fa:2:30:10 SLIDINGWINDOW:4:20 MINLEN:20). Clean reads were aligned to the *C. elegans* WBCel235 reference genome using BWA Mem (Li, 2013). PCR duplicated reads were removed using Picard tools. The Genome Analalysis toolkit (GATK) (Poplin et al., 2018) was then used to re-align the reads across variants using RealignerTargetCreator and IndelRealigner. Varscan (Koboldt et al., 2009) was used to identify differences (SNPs and Indels) between the samples and the reference genome. Finally, snpEff (Cingolani et al., 2012) was used to annotate the variants.

### RNA extraction and RNA-sequencing

Five freeze-thaw cycles were performed with Trizol samples that were stored at −70°C (see section ‘Collection of synchronized L1-staged *C. elegans’*), by thawing samples in a warm or hot water bath and immediately refreezing samples in liquid nitrogen. RNA extraction and transcriptome sequencing was performed by BGI Tech Solutions (HongKong) Co.Limited. RNA samples were sequenced using the BGI DNBSEQ Eukaryotic Strand-specific Transcriptome Resequencing product.

### RNA-sequencing analysis

RNA sequencing reads from the three experimental sets and the developmental time-course RNA-seq data (Meeuse et al., 2020) were treated in the same way. Raw reads were trimmed for adapters, low quality sequences and short reads with Trimmomatic (Bolger et al., 2014) (Version 0.39, parameters: ILLUMINACLIP: TruSeq3-SE.fa:2:30:10 SLIDINGWINDOW:4:20 MINLEN:20). Remaining ribosomal RNA was removed with sortmeRNA (Kopylova et al., 2012) (Version 2.1, default parameters), clean reads were aligned to the *C. elegans* WBCel235 reference genome and gene counts quantified with salmon (Patro et al., 2017) (Version 1.9.0, parameters: --gcBias, -- sepBias, -l A). Differentially expressed genes were then identified with DESeq2 (FDR < 0.01) (Love et al., 2014). Enrichment for previously published mutant RNASeqs was performed with worm Exp 1.0 (Yang et al., 2016). All visualisation was produced with the programming language R and the ggplot2 library.

#### Developmental progression estimation

Developmental progression was estimated by comparing gene expression data to a gene expression time-course obtained from Meeuse et al. (Meeuse et al., 2020) where hourly time points were taken from arrested and synchronized L1 larval animals and sequenced. Our data also came from arrested synchronized L1 animals that had been recovered for 6 hours. First, both our data and the Meeuse et al. data were aligned to the same reference to obtain read counts. Then our gene expression profile was compared to the time course profile by first using PCA to embed the Meeuse data alone. Using the eigenvectors obtained from this embedding, our data was embedded into the same space. From this the distance from each point to the piecewise linear curve determined by the time course data was computed. Finally, the time from the nearest two time points was interpolated and used as the estimated developmental time of our samples (see Figure S4B). Data was plotted using a custom function written in MATLAB for plotting raw data points with random jitter, along with the mean and 95% confidence interval.

### MNase-sequencing

#### In vivo MNase digestion

*In vivo* MNAse digestion was performed on purified nuclei isolated from *C. elegans* (wild type, *swsn-1* single, *swsn-1; snfc-5* double and *swsn-1; snfc-5; ubr-5* triple mutant) L1 frozen “worm balls” (see section ‘Collection of synchronized L1-staged *C. elegans*’). Prior to nuclei harvest, frozen “worm balls” containing 55,000 to 60,000 L1s for each strain were individually homogenized in a liquid nitrogen cryo cup. Nuclei purification was optimized using a nuclei pure prep nuclei isolation kit (NUC-201, Sigma Aldrich). Briefly, 400µl of ice-cold Lysis Solution containing DTT and Triton X-100 was added to each homogenized sample, vortexed and incubated on ice for 5 minutes. Cell lysis and nuclei morphology was determined by microscopic examination to ensure proper homogenization by taking 2µl sample. Nuclei were purified by centrifugation through 1.8 M sucrose cushion solution according to the manufacturer’s protocol. Briefly, 900µl of cold 1.8 M sucrose cushion solution was added to each 500µl lysate sample on ice and gently mixed. For each sample preparation, 500µl of ice cold 1.8 M sucrose cushion solution was added to the bottom of a fresh 2ml Eppendorf tube on ice. A total of 1.4ml of lysate solution from the previous step was then slowly layered on top of the 500µl of sucrose cushion solution and set for bench top centrifugation for 45 minutes at 13,000 rpm at 4°C. Supernatant (cytoplasm and cell debris) and the clear sucrose cushion layers were removed without disturbing the pellet of purified nuclei at the bottom of each tube. Nuclei pellet was vortexed briefly and resuspended with 500µl cold Nuclei PURE Storage Buffer. Nuclei pellet were collected by centrifugation at 500g for 5 minutes at 4°C, resuspended again with 50μl cold Nuclei PURE Storage Buffer and vortexed again to completely suspend the nuclei pellet. Qubit HS dsDNA quantification was performed on the purified nuclei to estimate the nucleic acid content. Purified Nuclei of 500ng/reaction were digested with MNAse (M0247S, NEB). The concentration of MNase was titrated for each reaction to obtain mononucleosomes. 250U/ml resulted in mononucleosomes for replicate 1. A concentration of 200U/ml resulted in mononucleosomes for replicate 2. Digestion was for 15 minutes at 37°C. MNase digestion was terminated by adding stop solution containing 3% SDS, 20mM EDTA (final concentrations). Mononuclesomes were treated with Proteinase K (Invitrogen) and incubated at 65°C for 1 hour followed by DNA purification using Zymo clean and Concentrator. Mononucleosome bands were confirmed by D5000 ScreenTape in an Agilent 2200 TapeStation system.

For library preparation, DNA was measured again with Qubit dsDNA HS Assay Kit and 65ng of mononucleosomal purified DNA was used for NEBNext Ultra II DNA Library Prep Kit (Illumina) with size selection. Nucleosomal DNA libraries were pooled in groups of 8 per lane and quantified with by TapeStation (Agilent 2200 TapeStation system) run using a D1000 ScreenTape (Agilent). Libraries were sequenced on an NextSeq 2000 machine (Illumina) in paired end mode with 60bp read length.

#### MNAse-seq data processing and analysis

The sequenced paired-end reads were mapped to the *C. elegans* (ce11) genome using bowtie2 aligner v.2.2.9 (Langmead and Salzberg, 2012) with default parameters. The resulting bam files were converted to bigwig tracks using deeptools bamCoverage (Ramírez et al., 2016) with parameters -bs 1 --extendReads 148 –normalizeUsing RPGC --effectiveGenomeSize 98259998. Additional paraments for bamCoverage was comprised of --MNase --minFragmentLength 100 -- maxFragmentLength 200 --Offset 1 2 and --smoothLength 30. Peak calling was performed with MACS2 (Zhang et al., 2008) with default parameters and reads were shifted by 75bp to cover the nucleosome dyad and extended to 150bp. Deeptools computeMatrix reference-point was used to compute the nucleosome density across a window spanning the TSS (500bp upstream to 700bp downstream). The TSS positions were extracted from Serizay et al. (Serizay et al., 2020), as the midpoint of the 300bp regions annotated as ”promoter”. Assignments of each promoter to Ubiquitous, Germline or Somatic was taken from these annotations. The resulting files were read in to R using the read.table function. To produce metagene plots spanning the region surrounding the TSS, the signal at 10bp intervals across the region was averaged across all promoters within the categories above. The ”background” intensity was defined as the minimum value across each trace and was subtracted from each position across the window. A clear signal indicating the expected nucleosome density was only observed for Ubiquitous genes, as expected since somatic promoters do not show such well-defined nucleosome densities (Serizay et al., 2020) and L1s do not have extensive germline tissue (Sulston and Horvitz, 1977). To compare the intensity of the +1 nucleosome across all ubiquitous promoters, the signal spanning from the TSS to 180bp downstream was extracted for each promoter and the signal for wild type subtracted from the mutant signal. To link changes in gene expression to changes in nucleosome density at promoters, the coordinates of the genes were extended by 500bp upstream and intersected with the 300bp promoter regions using bedtools slop and intersect respectively. Nine of the upregulated and 14 of the downregulated genes overlapped with ubiquitous promoters and the +1 signal was extracted and compared across strains.

### Generation of C. *elegans* protein extracts

Frozen L1 *C. elegans*/M9 pellets (see section ‘Collection of synchronized L1-staged *C. elegans*’) were thawed on ice, and 25µl DEPC H_2_O and 125µl of 2x protein lysis buffer (50mM Tris (pH 7.5), 300mM NaCl, 3mM MgCl_2_, 2mM dithiothreitol (DTT), 1% Igepal, complete proteinase inhibitor cocktail (Roche)) were added to the 100µl L1/M9 mix. Samples were transferred into 2ml Beadbug^TM^ tubes prefilled with 0.5mm zirconium beads (Merck Life Science) L1s were then broken up to extract proteins using a Beadbug homogenizer (Merck Life Science) at the highest setting for 90 seconds three times at 4°C. After every homogenisation step with the Beadbug, samples were centrifuged at maximum speed at 4°C for 2 minutes followed by 30 seconds at room temperature to remove bubbles and to check the lysis of the animals. Lysates were transferred into 1.5ml protein lobind tubes (Eppendorf), centrifuged at maximum speed for at least 30 minutes at 4°C and supernatants were transferred into new 1.5ml tubes. Subsequently, protein concentrations were determined using the Pierce BCA protein assay kit (Thermos Fisher Scientific) according to the manufacturer’s instructions and measuring the absorbance at 562nm with a spectrophotometer (Microplate Absorbance Reader, Hidex). Relative protein concentrations of samples were calculated using a bovine serum albumin (BSA) standard curve. Protein extracts were snap-frozen in liquid nitrogen and stored at stored at −70°C or used directly.

### SDS-PAGE

For sodium dodecyl sulphate polyacrylamide gel electrophoresis (SDS-PAGE), 25-30µg of *C. elegans* L1 protein extracts were supplemented with 10x NuPAGE Sample Reducing Agent and 4x LDS sample buffer (Thermo Fisher Scientific) to a final 1x concentration and denatured at 95°C for 5 minutes. Proteins were resolved alongside a protein ladder (PageRuler Plus Pre-stained Protein Ladder, Thermo Fisher Scientific) on NuPAGE 4-12% BisTris gradient gels using NuPAGE MOPS SDS running buffer (Thermo Fisher Scientific). Gel electrophoresis was routinely performed at 150V for approximately 1 hour and 15 minutes in a XCell SureLock electrophoresis system (Thermo Fisher Scientific) connected to a Bio-Rad PowerPac (Bio-Rad).

### Western blotting

Separated proteins were wet transferred onto 0.45mm pore-sized PVDF membranes (Immobilon®-FL PVDF membrane, Millipore) for 2 hours at 250mA and 4°C using cold transfer buffer (1.5x NuPAGE transfer buffer (Thermo Fisher Scientific), 10% methanol, 250µl 20% SDS/L) in a Mini Trans-Blot Cell (Bio-Rad). After the transfer, PVDF membranes were washed in water for 5 minutes, followed by a 2-minute shaking wash in methanol and another 5-minute wash in water. Membranes were air dried, rehydrated in methanol and TBST and incubated with Revert 700 Total Protein Stain (Licor) according to the manufacturer’s instructions. To obtain total protein signals for protein signal normalisation, membranes were imaged in a Licor Odyssey Imager in the 700nm channel.

Subsequently, membranes were blocked in 5% (w/v) milk in TBS-0.1% Tween 20 (TBST) for 1 hour at room temperature shaking, followed by an overnight 4°C rotating incubation with the mouse anti-FLAG M2 primary antibody (Simga Aldrich) at 1:1000 dilution in 5% milk. After three 5-minute shaking washes in TBST, membranes were incubated with a secondary IRDye 800CW anti-Mouse antibody (Licor) at 1:10,000 dilution in 5% milk in TBST for one hour at room temperature. Finally, membranes were washed again three times in TBST for 5 minutes shaking and imaged in the 800nm channel of the Licor Odyssey Imager to obtain SWSN-1::FLAG signals.

### Quantification of SWSN-1::FLAG protein levels

To quantify protein levels obtained by Western blotting, detected SWSN-1::FLAG signals were normalized to total protein signals. For this purpose, we subtracted the background of the 800nm SWSN-1::FLAG images by creating background images using the morphological opening (imopen) function with a disk shaped structuring element of radius 75 pixels and taking the absolute difference between the image and this background using MATLAB (v2021a Mathworks, Natick, MA). In Fiji (version 2.1.0/1.53c) (Schindelin et al., 2012), SWSN-1::FLAG signals of the background subtracted 800nm images were measured by recording the raw integrated densities within a rectangle drawn around the detected SWSN-1 band. This rectangle was moved across the image to record the SWSN-1::FLAG signals of every sample without changing its size. Furthermore, total protein signals of the 700nm images were recorded in Fiji, to determine the relative amounts of proteins that were loaded into each lane of the polyacrylamide gel and transferred onto the PVDF membranes. To do this, the raw total protein stain 700nm images were rotated to straighten the lanes if necessary. Then, a line with a width of 10 was drawn from top to bottom through one of the protein lanes covering all proteins. This line was moved across all sample lanes of the image and the line profile data for every lane was recorded from a central position of each lane. In excel, the wild-type SWSN-1::FLAG line profile data was plotted against each *swsn-1* single, *swsn-1; snfc-5* double and *swsn-1; snfc-5 ubr-5* triple mutant SWSN-1::FLAG line profiles and linear trendlines were added. The line profiles have intensity peaks where the bands occur in the total protein Western blot. The relative total protein levels were computed by comparing the intensity of each of these peaks in each of the lanes and determining a multiplicative factor that relates their relative intensities. To align the peaks to compute this factor, we performed a lag cross correlation correction to test if the collected line profiles matched well together or if shifting the individual data points up or down by a certain lag increased the correlation between the wild-type and mutant line profiles. We performed the lag cross correlation correction using xcorr function in MATLAB. The slope of linear trendlines for the best correlated lag were calculated and used as dilution factors to normalize the recorded mutant SWSN-1::FLAG signals to; the dilution factor for wild-type SWSN-1::FLAG is 1. The total protein normalized raw integrated densities of SWSN-1::FLAG signals of five independent quantitative Western blots were plotted in Prism (version 9.0.0) and a repeated measurements one-way ANOVA was performed to determine if samples were from the same distribution. After rejecting the null hypothesis, Tukey’s multiple comparison test was used to compare FLAG signals between all the conditions.

### Label-free proteomics and mass spec

#### Label-free proteomics

For proteome analysis, protein lysates (see section ‘Generation of *C. elegans* protein extracts’) were supplemented with LDS Sample Buffer (NuPAGE LDS sample buffer, Thermo Fisher Scientific) with 1 mM dithiothreitol (DTT). Samples were heated at 70°C for 10 min, alkylated by addition of 5.5 mM chloroacetamide, and 50 µg were loaded onto 4-12% gradient Bis–Tris gels. Proteins were separated by SDS–PAGE, stained using the Colloidal Blue Staining Kit (Life Technologies) and in-gel digested using trypsin. Peptides were extracted from gel and desalted on reversed-phase C18 StageTips (Rappsilber et al., 2007).

#### Mass spec analysis

Peptides were analyzed on a quadrupole Orbitrap mass spectrometer (Exploris 480, Thermo Scientific) equipped with a UHPLC system (EASY-nLC 1200, Thermo Scientific) as described (Bekker-Jensen et al., 2020; Kelstrup et al., 2012). The mass spectrometer was operated in data-dependent mode, automatically switching between MS and MS2 acquisition. Survey full scan MS spectra (m/z 300–1,700) were acquired in the Orbitrap. The 15 most intense ions were sequentially isolated and fragmented by higher energy C-trap dissociation (HCD) (Olsen et al., 2007). An ion selection threshold of 5,000 was used. Peptides with unassigned charge states, as well as with charge states < +2, were excluded from fragmentation. Fragment spectra were acquired in the Orbitrap mass analyzer.

#### Peptide identification

Raw data files were analyzed using MaxQuant software (development version 1.5.2.8) (Cox and Mann, 2008). Parent ion and MS2 spectra were searched against a database containing 28420 *C.elegans* protein sequences obtained from WormBase (WS269 release), as well as against 4190 proteins in the Ensembl Bacteria *E. coli* REL606 database using the Andromeda search engine (Cox et al., 2011). Spectra were searched with a mass tolerance of 6 ppm in MS mode, 20 ppm in HCD MS2 mode, strict trypsin specificity and allowing up to two miscleavages. Cysteine carbamidomethylation was set as a fixed modification, whilst N-terminal acetylation and oxidation were set as variable modifications. The dataset was filtered based on posterior error probability (PEP) to arrive at a FDR < 1% estimated using a target-decoy approach (Elias and Gygi, 2007). LFQ quantification was performed in the MaxQuant software using at least two LFQ ratio counts; the fast LFQ and the match between run options were turned on.

#### Data processing

Processed data from MaxQuant was analyzed in Perseus (version 1.6.15.0) (Tyanova et al., 2016) and visualized with RStudio (v. 4.1). Proteins or peptides flagged as “reverse”, “only identified by site” or “potential contaminant” were excluded from downstream analysis. Only proteins identified by no less than two peptides and at least one unique peptide were used in downstream analysis. Proteins mapped to the *E.coli* protein database were discarded. This dataset was further filtered so that proteins identified in at least two out of four replicates for each condition were kept. Missing values were imputed using random values from a Gaussian distribution using the default parameters in Perseus. P-values were calculated with a standard two-sided t-test and Benjamini-Hochberg correction was used for FDR calculation. Data visualization was performed using in-house R scripts and with existing libraries (ggplot2-v 3.3.5, ggrepel-v 0.9.1, RColorBrewer-v 1.1.2, VennDiagram-v 1.7.1) (see Figure S3F-G). Statistical analyses and plotting of data for Figure 3F were conducted using GraphPad Prism (v. 9.0.0).

#### Linear regression analysis of SWI/SNF protein levels

To determine the effect of the *swsn-1; snfc-5* double and *swsn-1; snfc-5; ubr-5* triple mutants on the protein levels of the SWI/SNF complex as a whole, a linear regression model was fit using the fitlm function in MATLAB (v2021a, Natick MA). The model was fit using both the subunit/protein identity and the genetic background as independent variables. As these are non-numeric variables, the fitlm procedure assigns an arbitrary value to obtain the best linear fit (see Figure S4B-E). fitlm uses an ANOVA to return an f-statistic which tests the null hypothesis that the model is not different from a null model, and also provides a t-statistic for the null hypothesis that each coefficient is different from 0. This procedure then estimates the best fit coefficient for each subunit (this can be thought of as the correction factor for the varying subunit expression levels) and regresses this out, leaving the effect of the genetic background (see Figure 4H). For reference, randomly selected sets of 12 proteins were analyzed in the same way, first each set was fit with a linear regression model and then the mean adjusted LFQ intensity was calculated for each of the genetic backgrounds. 100 such bootstrap samples were generated and their adjusted mean expression levels are shown as black dots with grey lines indicating the standard deviation in Figure 4H. These data were plotted using a custom function in MATLAB to plot the adjusted response.

## QUANTIFICATION AND STATISTICAL ANALYSIS

Statistical analyses are described in the individual methods sections. Furthermore, figure legends specify what statistical test and p-value cut-off was used to determine statistical significance.

## Acknowledgements

We thank the Gurdon Institute Media Kitchen for their support in providing reagents and media. We thank Kay Harnish for his support in managing the Gurdon Institute Sequencing Facility. We are grateful for the Miska Laboratory members, especially Giulia Furlan and Miguel Almeida, and Yaron Galanty for helpful discussions, and Marc Ridyard for laboratory management and maintenance of our nematode collection. Some *C. elegans* strains were provided by the CGC, which is funded by NIH Office of Research Infrastructure Programs (P40 OD010440). This work was supported by grants to E.A.M. from Cancer Research UK (C13474/A27826) and the Wellcome Trust (219475/Z/19/Z). Work in the Sarkies Laboratory was funded by the Medical Research Council (Epigenetics and Evolution), University of Oxford Department of Biochemistry and Lincoln College Oxford. This work was also supported by research grants to D.Q.M. from the National Institute of General Medical Sciences (NIGMS) [R01GM121597]. Work in the Beli Laboratory was funded by the Deutsche Forschungsgemeinschaft (DFG, German Research Foundation) - Project-ID 259130777 -SFB 1177. L.L. was supported by a Boehringer Ingelheim Fonds PhD fellowship.

## KEY RESOURCE TABLE

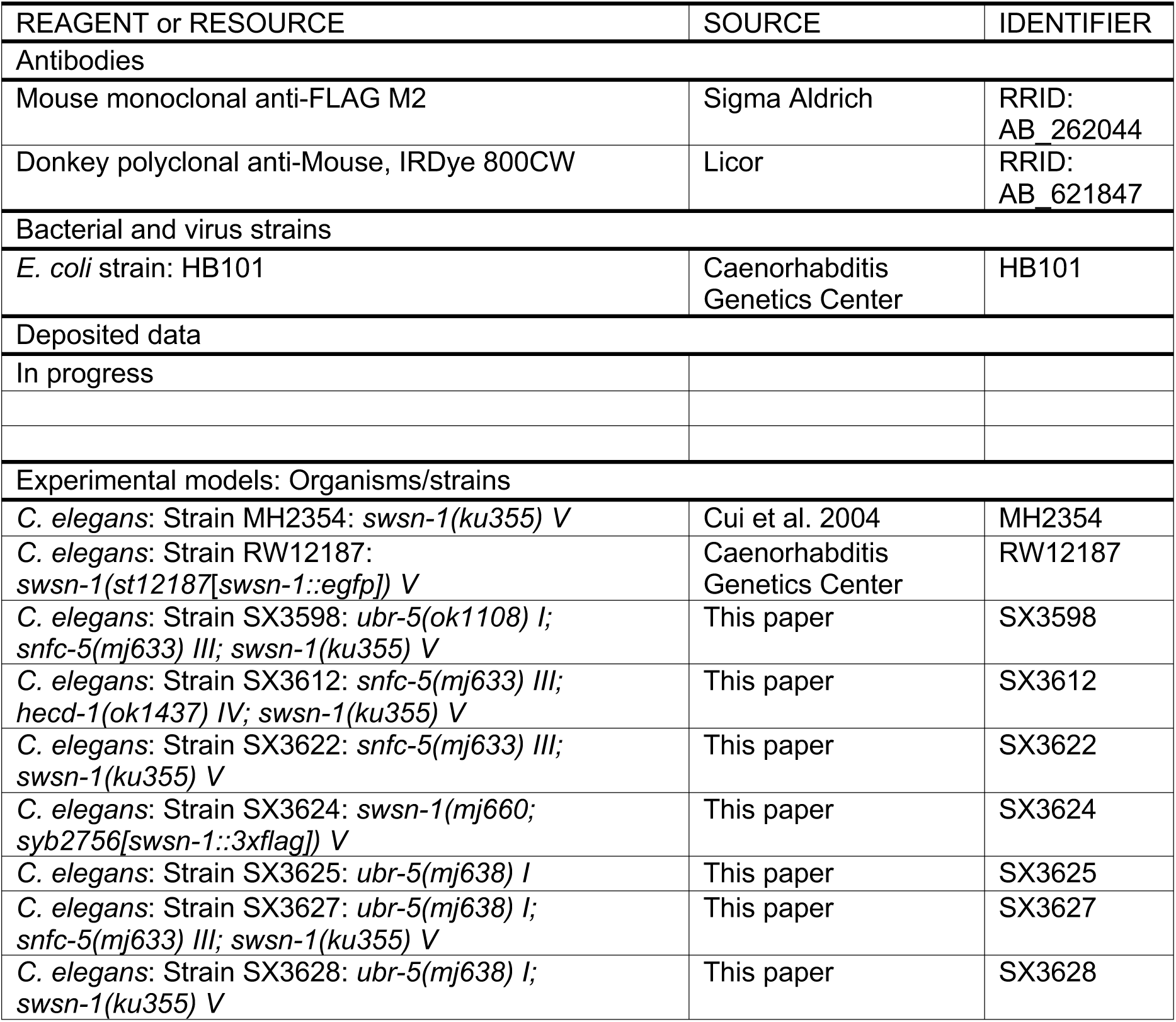
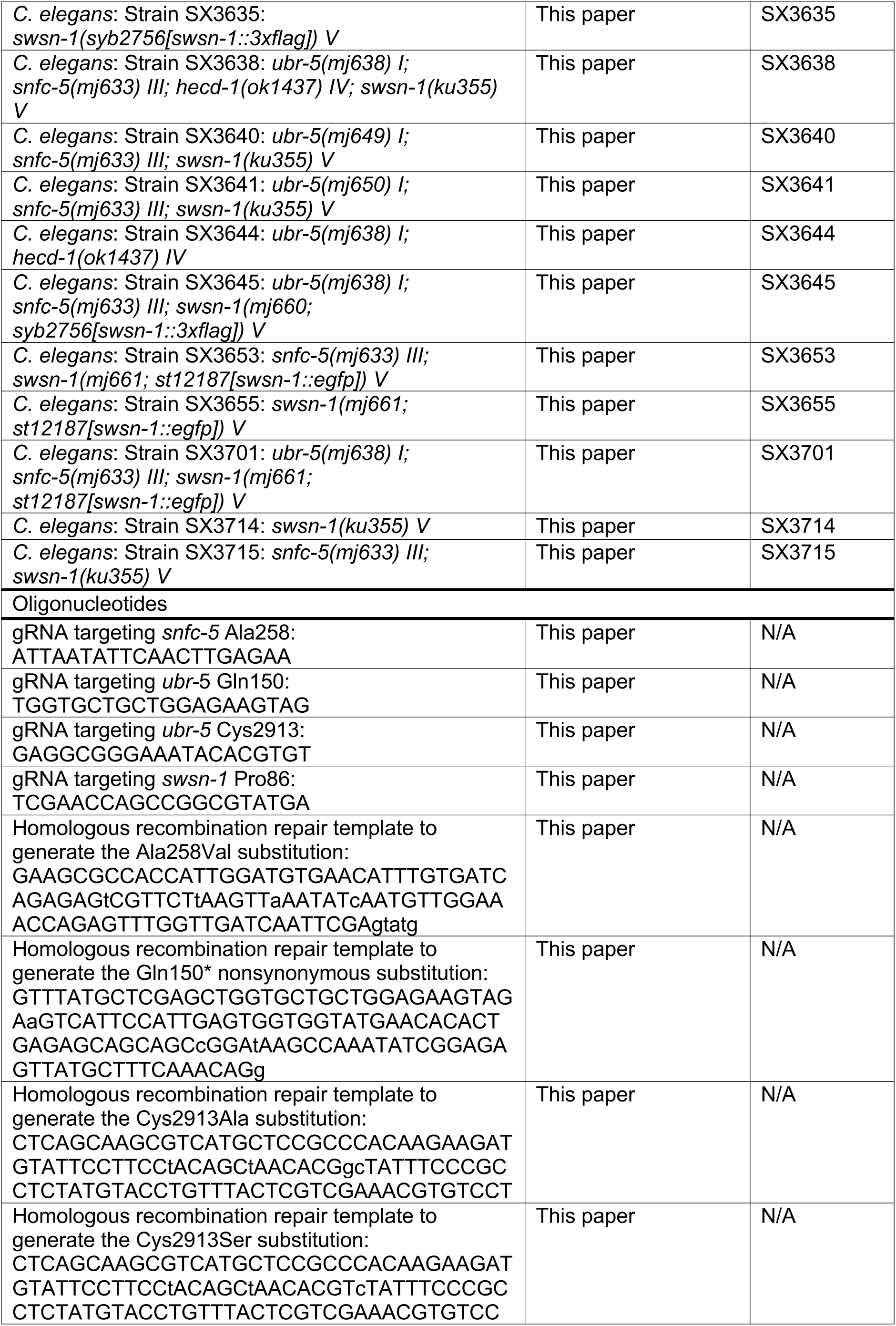
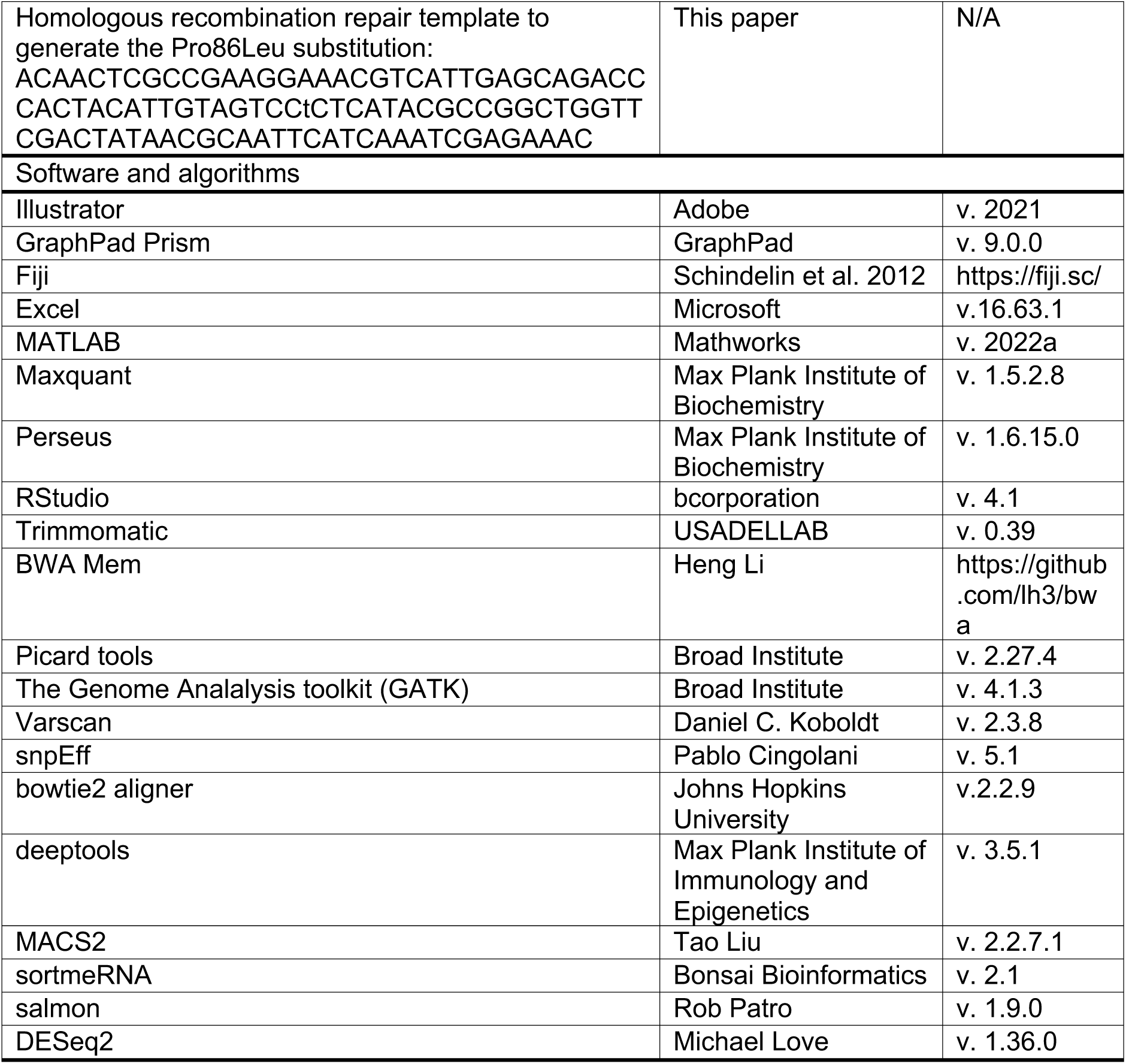

---

**Figure S1: Related to Figure 1.**
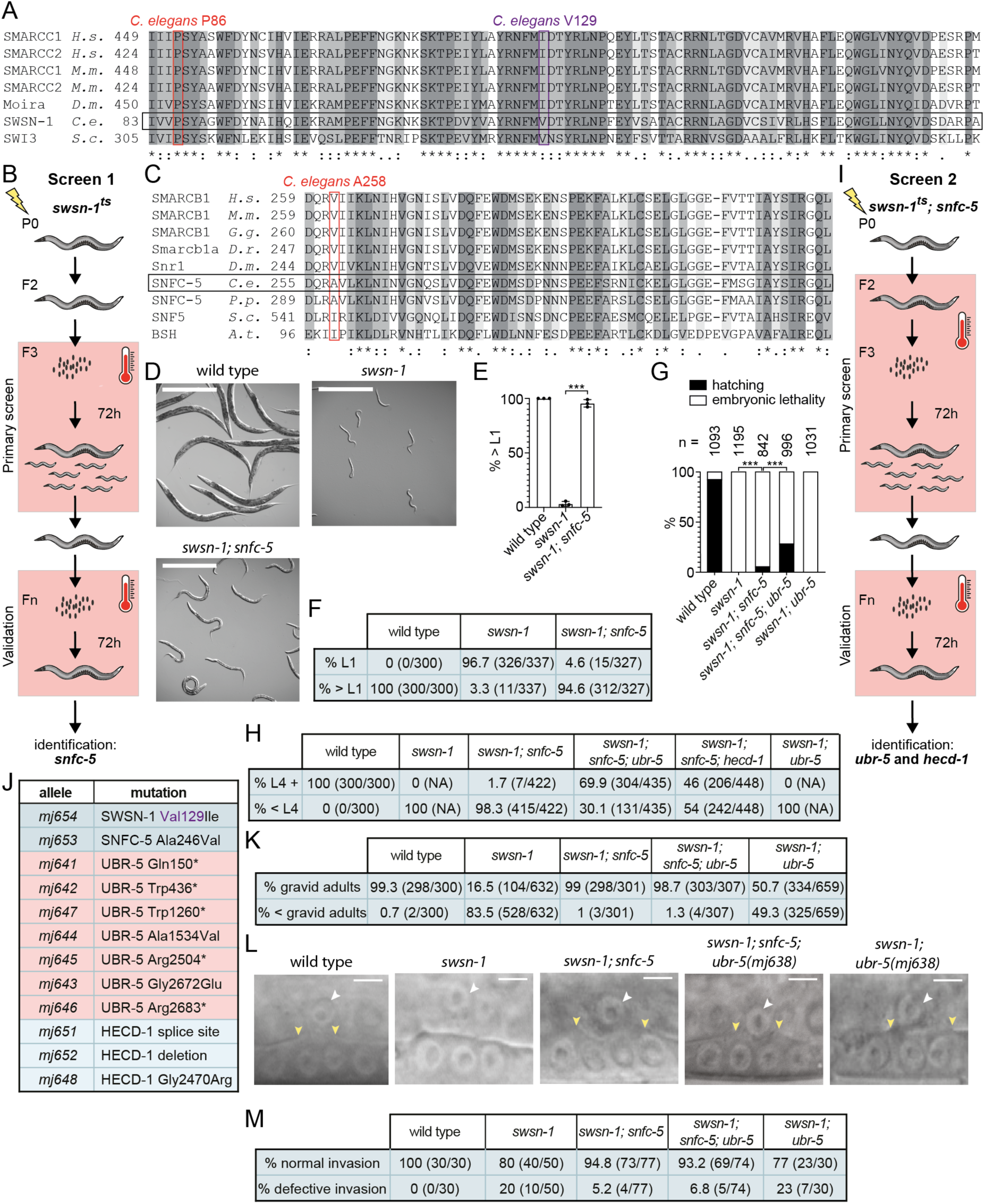
**A)** Alignments of the SWIRM domains (based on UniProt domain annotations) of human (*H.s.*) SMARCC1 and SMARCC2, mouse (*M.m.*) SMARCC1 and SMARCC2, fruit fly (*D.m.*) Moira, *C. elegans* (*C.e.*) SWSN-1 and yeast (*S.c.*) SWI3 using the UniProt protein alignment tool. ’*’ and dark grey shading = fully conserved residues, ’:’ and grey shading = strongly similar residues (> 0.5 in the Gonnet PAM 250 matrix) and ’.’ and light grey shading = weakly similar residues (< 0.5). The conserved proline mutated in *swsn-1(ku355)* animals is highlighted by a red rectangle. The valine that is mutated in a mutant from the second screen (see J) is highlighted by a purple rectangle. **B)** Schematic of the EMS mutagenesis screen with *swsn-1* temperature-sensitive (ts) mutants. The restrictive temperature of 25°C is indicated by the red box. **C)** Alignments of the repeat 2 domain (based on UniProt domain annotation of SMARCB1) of *H.s.* SMARCB1, *M.m.* SMARCB1, chicken (*G.g.*) SMARCB1, zebrafish (*D.r.*) Smarcb1a, *D.m.* Snr1, *C.e.* SNFC-5, *Pristionchus pacificus* (*P.p.*) SNFC-5, *S.c.* SNF5 and Arabidopsis thaliana (*A.t.*) BSH using the UniProt protein alignment tool. Similar and conserved residues are highlighted as described in A. The alanine mutated in *C.e. snfc-5* mutants identified in the first screen is highlight by a red rectangle. **D-F)** Quantification of *C. elegans* developmental stages after exposing embryos to 25°C for 48 hours. (D) Representative images of wild-type animals compared to *swsn-1* single and *swsn-1; snfc-5* double mutants, scale bar = 500μm. (E) Percentage of animals older than the larval 1 (L1) stage of >=100 scored animals (n=3). Bar heights represent the mean, error bars represent standard deviation, *** = Bonferroni corrected Fisher’s exact test p-value < 0.0001. (F) Contingency table containing combined developmental stage scorings from triplicates of E. This table was used for Fisher’s exact test p-value calculations. **G)** Quantification of larval hatching or embryonic lethality at 25°C 24 hours after collecting 842-1195 embryos from wild-type animals, *swsn-1* single, *swsn-1; snfc-5* double; *swsn-1; snfc-5; ubr-5* triple and *swsn-1; ubr-5* double mutants that were grown at 25°C for 16 hours before collecting the embryos. *** = Fisher’s exact test p-value < 0.0001. **H)** Contingency table containing combined developmental stage scorings from triplicates of Figure 1B. This table was used for Fisher’s exact test p-value calculations. *swsn-1* single and *swsn-1*; *ubr-5* double mutants were not scored, because no larvae hatched. **I)** Schematic of the EMS mutagenesis screen with *swsn-1; snfc-5* mutants using more stringent conditions than in B. The restrictive temperature of 25°C is indicated by the red box. **J)** Table of potential causative mutations of mutants recovered in I. **K)** Contingency table containing combined developmental stage scorings from triplicates of Figure 1E. This table was used for Fisher’s exact test p-value calculations. **L-M)** Quantification of anchor cell (AC) invasion. (L) Representative images of wild-type animals (normal invasion) compared to *swsn-1* single (defective invasion), *swsn-1; snfc-5* double (normal invasion), *swsn-1; snfc-5; ubr-5* triple (normal invasion) and *swsn-1; ubr-5* double (normal AC invasion, despite defects observed in 23% of animals) mutants, ACs are indicated by white arrowheads, boundaries of breach in the BM are indicated by yellow arrowheads, scale bar = 5µm. (M) Contingency table containing AC invasion scorings of Figure 1G. This table was used for Fisher’s exact test p- value calculations. Alleles used: *swsn-1(ku355), snfc-5(mj633), ubr-5(mj638), hecd-1(ok1437)*.

**Figure S2: Related to Figure 2.**
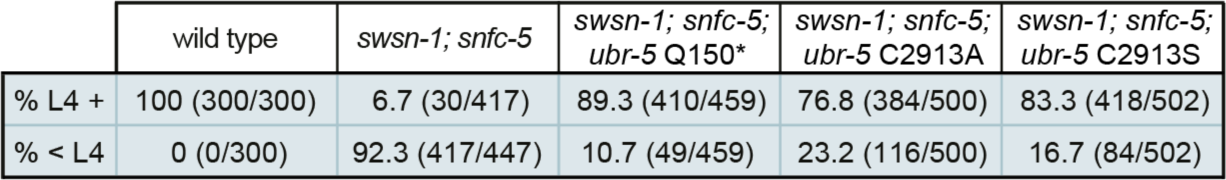
Contingency table containing combined developmental stage scorings from triplicates of Figure 2C. This table was used for Fisher’s exact test p-value calculations. Alleles used: *swsn-1(ku355), snfc-5(mj633), ubr-5(mj638), ubr-5(mj650), ubr-5(mj649)*

**Figure S3: Related to Figure 3.**
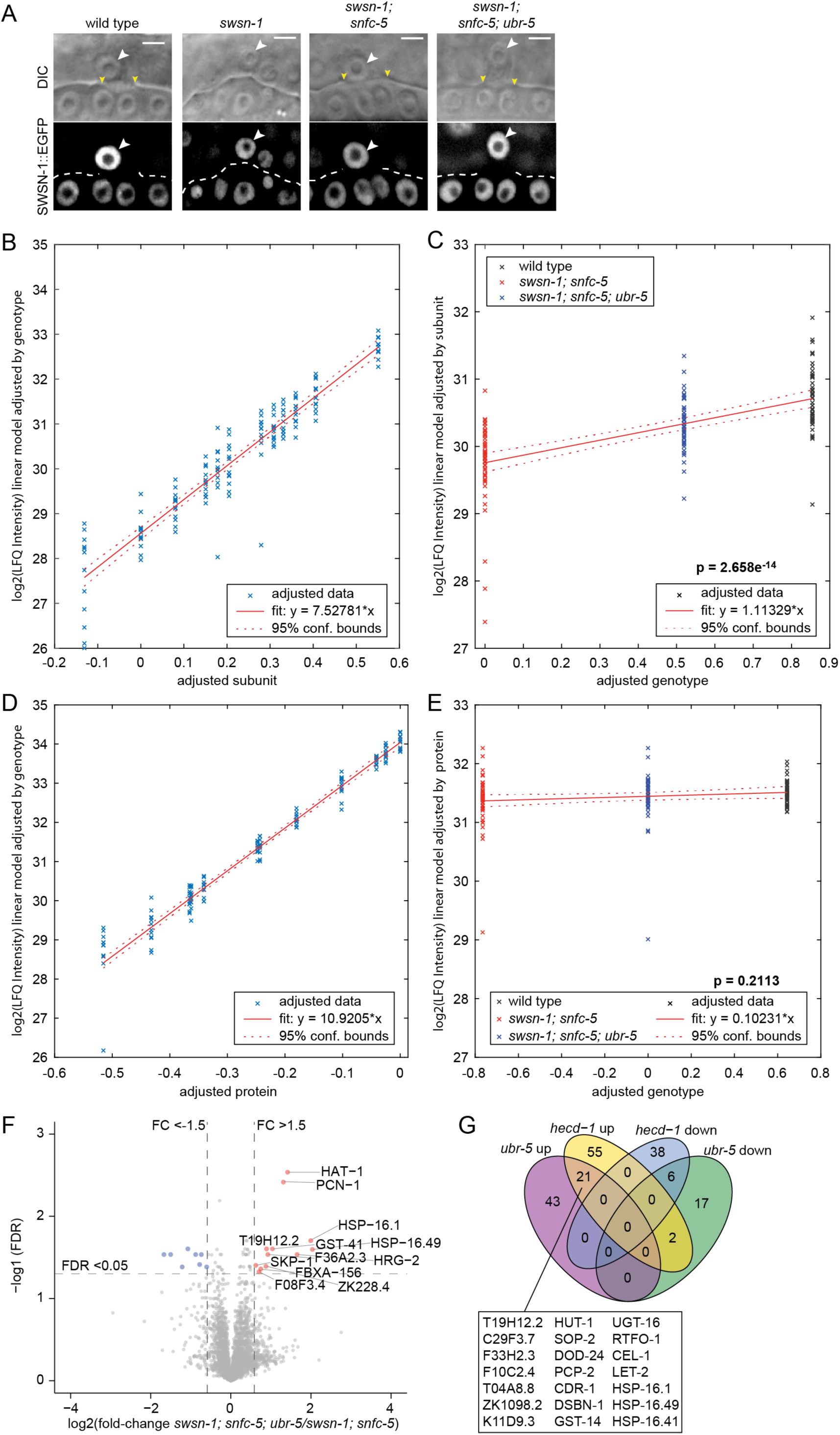
**A)** Quantification of SWSN-1::EGFP intensities in the AC of P6.p 4-cell-staged wild-type (*swsn-1::egfp*), *swsn-1* single mutant (*swsn-1^ts^::egfp)*, *swsn-1; snfc-5* double mutant *(swsn-1^ts^::egfp; snfc-5)* and *swsn-1; snfc-5; ubr-5* triple mutant *(swsn-1^ts^::egfp; snfc-5; ubr-5)* animals exposed to 25°C from the L2-L3 molt/early L3 stage until the P6.p 4-cell stage. Representative images of quantified animals, ACs are indicated by white arrowheads, boundaries of breach in the BM are indicated by yellow arrowheads, scale bar = 5µm. (The SWSN-1::EGFP images are also displayed in Figure 3D). **B-C)** Linear regression analysis of protein levels of the SWI/SNF subunits (B) Added variable plot for protein subunits displaying adjusted protein levels data (x) from Figure 3G of the three genotypes combined. The x-axis represents scaled distances of subunits to adapt their individual adjusted log2(LFQ intensities) to a linear fit (red line). The y-axis represents log2(LFQ intensities) of the linear model adjusted by genotype. (C) Added variable plot for genotypes displaying adjusted protein levels data (x) from Figure 3G of the twelve protein subunits combined. The x-axis represents scaled distances of genotypes to adapt their individual adjusted log2(LFQ intensities) to a linear fit (red line). The y-axis represents log2(LFQ intensities) of the linear model adjusted by subunit. ANOVA (p-value = 2.658e^-14^) was used to determine that the genotype has an effect on the protein levels of the set of SWI/SNF subunits. **D-E)** Linear regression analysis of protein levels of twelve randomly selected proteins (ARX-6, EXOS-1, CYN-1, NBET-1, EMB-4, ZK1236.5, VPS-29, TFTC-3, MDT-9, TBA-1, HMT-1 and Y39G8B.1) in synchronized L1-staged wild-type, *swsn-1; snfc-5* double mutant and *swsn-1; snfc-5 ubr-5* triple mutant animals determined by label-free proteomics mass spec quantification (n=4). (D) Added variable plot for the twelve proteins displaying adjusted protein levels data of the three genotypes combined. The x-axis represents scaled distances of the proteins to adapt their individual adjusted log2(LFQ intensities) to a linear fit (red line). The y-axis represents log2(LFQ intensities) of the linear model adjusted by genotype. (E) Added variable plot for genotypes displaying adjusted protein levels data of the twelve proteins combined. The x-axis represents scaled distances of genotypes to adapt their individual adjusted log2(LFQ intensities) to a linear fit (red line). The y-axis represents log2(LFQ intensities) of the linear model adjusted by protein. ANOVA (p-value = 0.02113) was used to determine that the genotype has no effect on the levels of this set of twelve randomly selected proteins. **F)** Volcano plot of up- and downregulated proteins in *swsn-1; snfc-5; ubr-5* triple mutants versus *swsn-1; snfc-5* double mutants determined by mass spec using label-free quantification. Fold change < −1.5 or > 1.5, FDR < 0.05. (n=4). **G)** Venn diagram of the overlap between up- and down-regulated proteins in *ubr-5* single and *hecd-1* single mutants compared to wild type animals determined by mass spec using label-free quantification. Fold change < −1.5 or > 1.5, two-sided t-test p-value < 0.05 (n=4). Alleles used: *swsn-1(ku355), swsn-1(syb2756[swsn-1::3xflag]), swsn-1(mj660; syb2756[swsn-1::3xflag]), swsn-1(st12187[swsn-1::egfp]), swsn-1(mj661; st12187[swsn-1::egfp]), snfc-5(mj633), ubr-5(mj638)*.

**Figure S4: Related to Figure 4.**
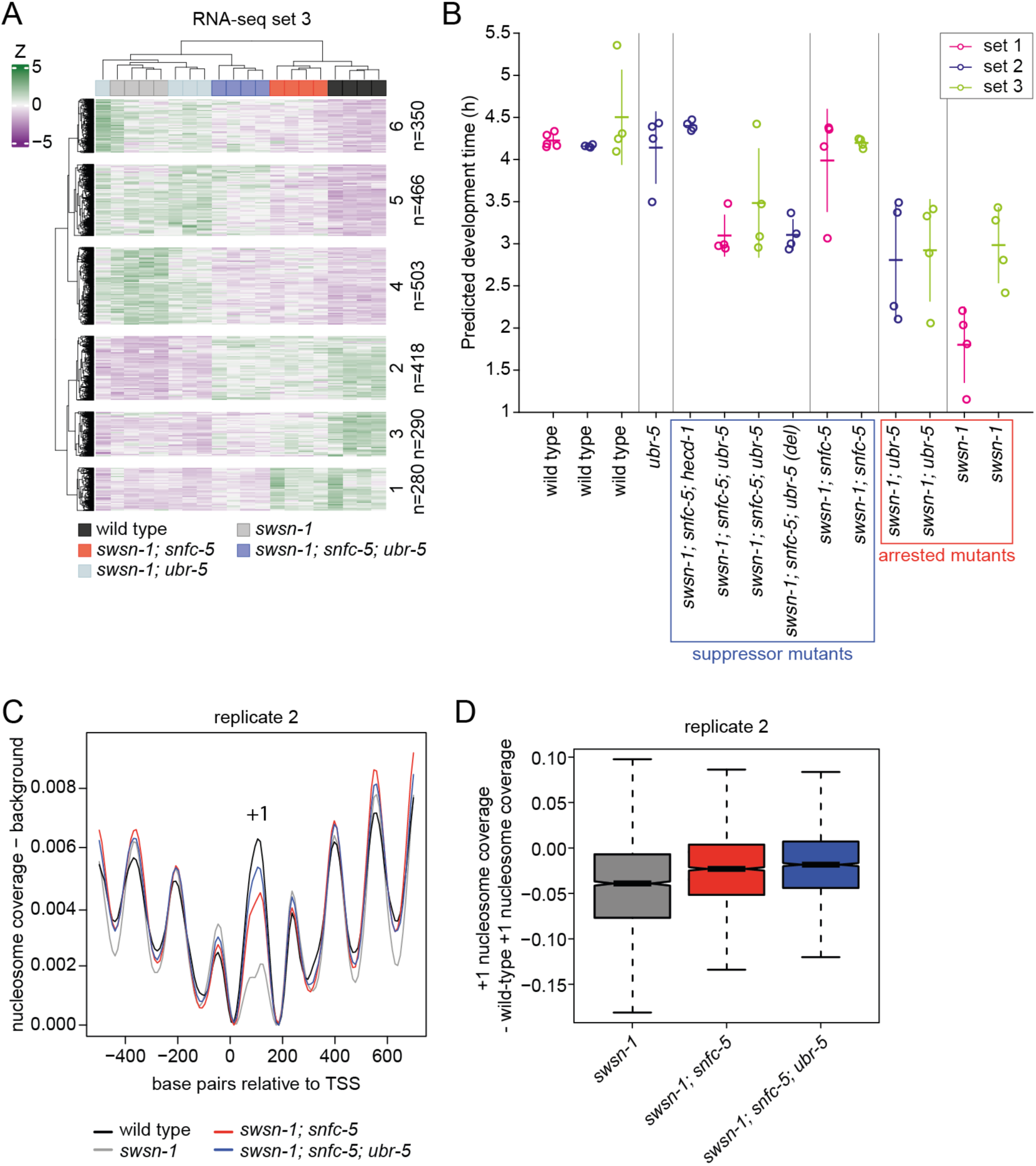
**A)** Z-score heatmap of the 2307 DEGs differentially expressed between wild type and at least one mutant from RNA-seq set 3 after K-means clustering. **B)** Predicted developmental progression of the L1-staged mutants and wild-type animals used for RNA-sequencing estimated by comparing their gene expression profiles to a published wild-type *C. elegans* time course (hourly from 1-24 hours of larval development) RNA-seq data set (Meeuse et al., 2020). Vertical lines represent the 95% confidentiality intervals and horizontal lines the mean. **C)** Nucleosome traces around the TSS of ubiquitous genes determined by MNase-seq of synchronized L1-staged wild-type animals and *swsn-1* single, *swsn-1; snfc-5* double and *swsn-1; snfc-5; ubr-5* triple mutants. **D)** Box plots of locus-by-locus + 1 nucleosome coverage of ubiquitous genes determined by MNase- seq in *swsn-1* single, *swsn-1; snfc-5* double and *swsn-1; snfc-5; ubr-5* triple mutants relative to wild-type coverage. Bold horizontal lines represent the median, boxes represent interquartile range and whiskers extend to the greatest point <=1.5 times the interquartile range. Alleles used: *swsn-1(ku355), snfc-5(mj633), ubr-5(mj638), ubr-5(ok1108)* (used in *swsn-1; snfc-5; ubr-5* triple mutant from RNA-seq set 2, indicated by ‘(del)’)*, hecd-1(ok1437)*.

**Table S1:** Fold-change of SWI/SNF protein levels in *ubr-5* mutant versus wild-type. FDR = false discovery rate, FDR < 0.05 highlighted in bold.

| Protein/subunit | FC *ubr-5* mutant versus wild-type | FDR |
| --- | --- | --- |
| SWSN-1 | 1.01 | 0.853 |
| SWSN-4 | 1.06 | 0.257 |
| SNFC-5 | 1.13 | 0.157 |
| SWSN-2.1 | 1.15 | 0.15 |
| SWSN-2.2 | 1.03 | 0.329 |
| SWSN-3 | 1.07 | 0.205 |
| SWSN-6 | 0.99 | 0.901 |
| PBRM-1 | 1.06 | 0.656 |
| **SWSN-7** | **1.11** | **0.037** |
| SWSN-9 | 1.09 | 0.49 |
| **PHF-10** | **1.23** | **0.035** |
| LET-526 | 0.94 | 0.303 |
| DPFF-1 | 1.0 | 0.982 |

**Table S2:**
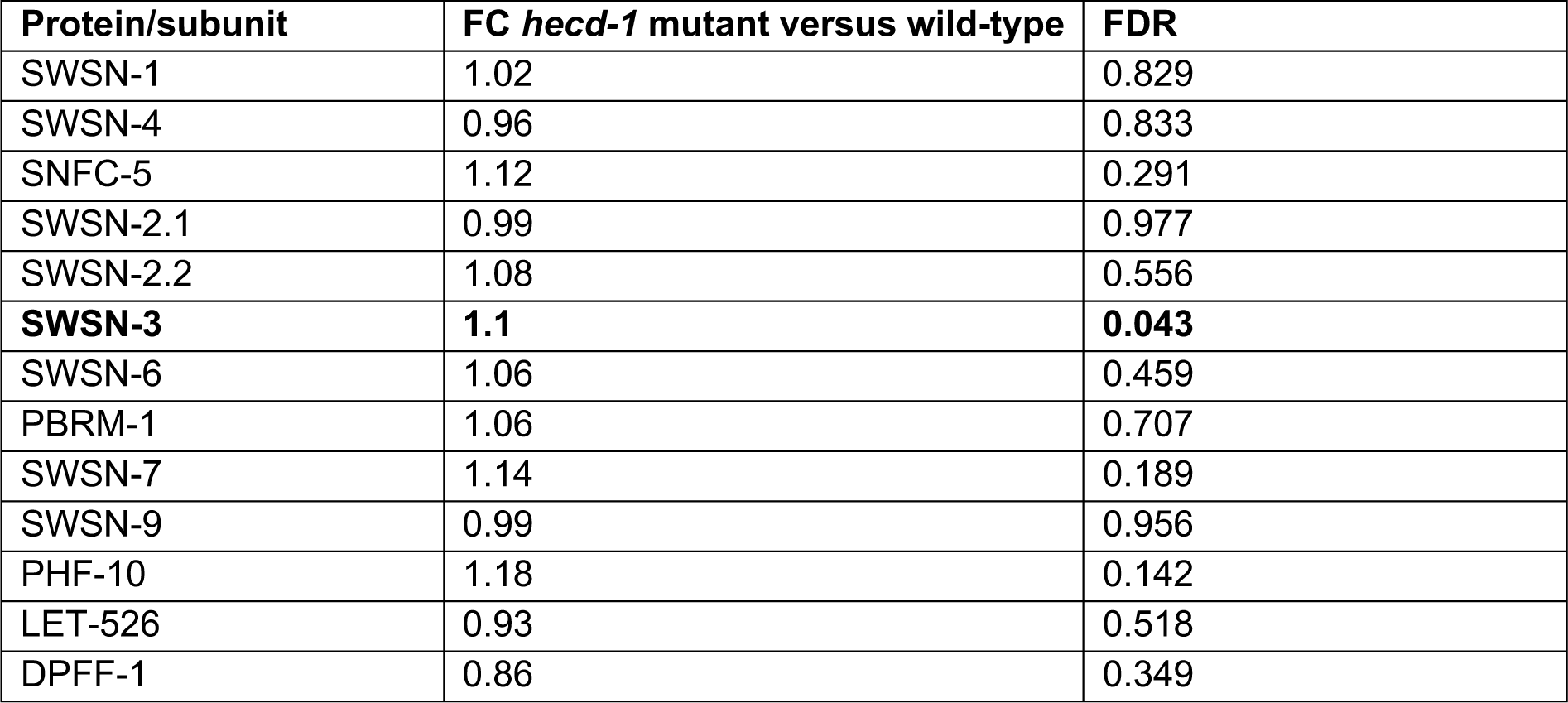
Fold-change of SWI/SNF protein levels in *hecd-1* mutant versus wild-type. FDR = false discovery rate, FDR < 0.05 highlighted in bold.

## Notes

### Competing Interest Statement

The authors have declared no competing interest.

## References

Akutsu, M., Dikic, I., and Bremm, A. (2016). Ubiquitin chain diversity at a glance. J. Cell Sci. 129, 875–880. https://doi.org/10.1242/JCS.183954/260238/AM/UBIQUITIN-CHAIN-DIVERSITY-AT-A-GLANCE.

Bekker-Jensen, D.B., Martínez-Val, A., Steigerwald, S., Rüther, P., Fort, K.L., Arrey, T.N., Harder, A., Makarov, A., and Olsen, J. V. (2020). A compact quadrupole-orbitrap mass spectrometer with FAIMS interface improves proteome coverage in short LC gradients. Mol. Cell. Proteomics 19, 716–729. https://doi.org/10.1074/mcp.TIR119.001906.

Bolger, A.M., Lohse, M., and Usadel, B. (2014). Trimmomatic: a flexible trimmer for Illumina sequence data. Bioinformatics 30, 2114–2120. https://doi.org/10.1093/BIOINFORMATICS/BTU170.

Brenner, S. (1974). The Genetics of Caenorhabditis elegans. Genetics 71–94. .

Bultman, S., Gebuhr, T., Yee, D., La Mantia, C., Nicholson, J., Gilliam, A., Randazzo, F., Metzger, D., Chambon, P., Crabtree, G., et al. (2000). A Brg1 null mutation in the mouse reveals functional differences among mammalian SWI/SNF complexes. Mol. Cell 6, 1287–1295. https://doi.org/10.1016/S1097-2765(00)00127-1.

Cingolani, P., Platts, A., Wang, L.L., Coon, M., Nguyen, T., Wang, L., Land, S.J., Lu, X., and Ruden, D.M. (2012). A program for annotating and predicting the effects of single nucleotide polymorphisms, SnpEff: SNPs in the genome of Drosophila melanogaster strain w1118; iso-2; iso-3. Fly (Austin). 6, 80–92. https://doi.org/10.4161/FLY.19695/SUPPL_FILE/KFLY_A_10919695_SM0001.ZIP.

Cox, J., and Mann, M. (2008). MaxQuant enables high peptide identification rates, individualized p.p.b.-range mass accuracies and proteome-wide protein quantification. Nat. Biotechnol. 26, 1367– 1372. https://doi.org/10.1038/NBT.1511.

Cox, J., Neuhauser, N., Michalski, A., Scheltema, R.A., Olsen, J. V., and Mann, M. (2011). Andromeda: a peptide search engine integrated into the MaxQuant environment. J. Proteome Res. 10, 1794–1805. https://doi.org/10.1021/PR101065J.

Cui, K., and Zhao, K. (2012). Genome-wide approaches to determining nucleosome occupancy in metazoans using MNase-Seq. Methods Mol. Biol. 833, 413–419. https://doi.org/10.1007/978-1-61779-477-3_24/COVER.

Cui, M., Fay, D.S., and Han, M. (2004). lin-35/Rb Cooperates With the SWI/SNF Complex to Control Caenorhabditis elegans Larval Development. Genetics 35, 1177–1185. https://doi.org/10.1534/genetics.103.024554.

Elias, J.E., and Gygi, S.P. (2007). Target-decoy search strategy for increased confidence in large-scale protein identifications by mass spectrometry. Nat. Methods 2007 43 *4*, 207–214. https://doi.org/10.1038/nmeth1019.

Euskirchen, G.M., Auerbach, R.K., Davidov, E., Gianoulis, T.A., Zhong, G., Rozowsky, J., Bhardwaj, N., Gerstein, M.B., and Snyder, M. (2011). Diverse roles and interactions of the SWI/SNF chromatin remodeling complex revealed using global approaches. PLoS Genet. 7. https://doi.org/10.1371/journal.pgen.1002008.

Flaus, A., Martin, D.M.A., Barton, G.J., and Owen-Hughes, T. (2006). Identification of multiple distinct Snf2 subfamilies with conserved structural motifs. Nucleic Acids Res. 34, 2887–2905. https://doi.org/10.1093/NAR/GKL295.

Frøkjær-Jensen, C., Wayne Davis, M., Hopkins, C.E., Newman, B.J., Thummel, J.M., Olesen, S.P., Grunnet, M., and Jorgensen, E.M. (2008). Single-copy insertion of transgenes in Caenorhabditis elegans. Nat. Genet. 40, 1375–1383. https://doi.org/10.1038/ng.248.

Garcia-Barcena, C., Osinalde, N., Ramirez, J., and Mayor, U. (2020). How to Inactivate Human Ubiquitin E3 Ligases by Mutation. Front. Cell Dev. Biol. 8, 39. https://doi.org/10.3389/fcell.2020.00039.

Hamada, F.N., Rosenzweig, M., Kang, K., Pulver, S.R., Ghezzi, A., Jegla, T.J., and Garrity, P.A. (2008). An internal thermal sensor controlling temperature preference in Drosophila. Nature 454, 217–220. https://doi.org/10.1038/nature07001.

Han, Y., Reyes, A.A., Malik, S., and He, Y. (2020). Cryo-EM structure of SWI/SNF complex bound to a nucleosome. Nature 579, 452–455. https://doi.org/10.1038/s41586-020-2087-1.

He, S., Wu, Z., Tian, Y., Yu, Z., Yu, J., Wang, X., Li, J., Liu, B., and Xu, Y. (2020). Structure of nucleosome-bound human BAF complex. Science (80-.). eaaz9761. https://doi.org/10.1126/science.aaz9761.

Kadoch, C., and Crabtree, G.R. (2015). Mammalian SWI/SNF chromatin remodeling complexes and cancer: Mechanistic insights gained from human genomics. Sci. Adv. 1. https://doi.org/10.1126/SCIADV.1500447/ASSET/4396DAC5-0BA9-49F8-8DA2-96264B0FC7DE/ASSETS/GRAPHIC/1500447-F6.JPEG.

Kelstrup, C.D., Young, C., Lavallee, R., Nielsen, M.L., and Olsen, J. V. (2012). Optimized fast and sensitive acquisition methods for shotgun proteomics on a quadrupole orbitrap mass spectrometer. J. Proteome Res. 11, 3487–3497. https://doi.org/10.1021/PR3000249/SUPPL_FILE/PR3000249_SI_001.ZIP.

Koboldt, D.C., Chen, K., Wylie, T., Larson, D.E., McLellan, M.D., Mardis, E.R., Weinstock, G.M., Wilson, R.K., and Ding, L. (2009). VarScan: variant detection in massively parallel sequencing of individual and pooled samples. Bioinformatics 25, 2283–2285. https://doi.org/10.1093/BIOINFORMATICS/BTP373.

Kopylova, E., Noé, L., and Touzet, H. (2012). SortMeRNA: fast and accurate filtering of ribosomal RNAs in metatranscriptomic data. Bioinformatics 28, 3211–3217. https://doi.org/10.1093/BIOINFORMATICS/BTS611.

Langmead, B., and Salzberg, S.L. (2012). Fast gapped-read alignment with Bowtie 2. Nat. Methods 2012 94 *9*, 357–359. https://doi.org/10.1038/nmeth.1923.

Large, E.E., and Mathies, L.D. (2014). Caenorhabditis elegans SWI/SNF Subunits Control Sequential Developmental Stages in the Somatic Gonad. G3 Genes, Genomes, Genet. 4, 471–483. https://doi.org/10.1534/g3.113.009852.

Li, H. (2013). Aligning sequence reads, clone sequences and assembly contigs with BWA-MEM. https://doi.org/10.48550/arxiv.1303.3997.

Love, M.I., Anders, S., and Huber, W. (2014). Differential analysis of count data - the DESeq2 package. Genome Biol. 15, 550. https://doi.org/110.1186/s13059-014-0550-8.

Mani, U., S, A.S., Goutham R N, A., and Mohan S, S. (2017). SWI/SNF Infobase—An exclusive information portal for SWI/SNF remodeling complex subunits. PLoS One 12. https://doi.org/10.1371/JOURNAL.PONE.0184445.

Mashtalir, N., D’Avino, A.R., Michel, B.C., Luo, J., Pan, J., Otto, J.E., Zullow, H.J., McKenzie, Z.M., Kubiak, R.L., St. Pierre, R., et al. (2018). Modular Organization and Assembly of SWI/SNF Family Chromatin Remodeling Complexes. Cell 175, 1272–1288.e20. https://doi.org/10.1016/J.CELL.2018.09.032.

Mathies, L.D., Lindsay, J.H., Handal, A.P., Blackwell, G.M.G., Davies, A.G., and Bettinger, J.C. (2020). SWI/SNF complexes act through CBP-1 histone acetyltransferase to regulate acute functional tolerance to alcohol. BMC Genomics 21, 1–15. https://doi.org/10.1186/s12864-020-07059-y.

Meeuse, M.W., Hauser, Y.P., Moya, L.J.M., Hendriks, G.-J., Eglinger, J., Bogaarts, G., Tsiairis, C., and Großhans, H. (2020). Developmental function and state transitions of a gene expression oscillator in Caenorhabditis elegans. Mol. Syst. Biol. 16, e9498. https://doi.org/10.15252/MSB.20209498.

Narayanan, R., Pirouz, M., Kerimoglu, C., Pham, L., Wagener, R.J., Kiszka, K.A., Rosenbusch, J., Seong, R.H., Kessel, M., Fischer, A., et al. (2015). Loss of BAF (mSWI/SNF) Complexes Causes Global Transcriptional and Chromatin State Changes in Forebrain Development. Cell Rep. 13, 1842– 1854. https://doi.org/10.1016/J.CELREP.2015.10.046.

Neigeborn, L., and Carlson, M. (1984). Genes Affecting the Regulation of SUC2 Gene Expression by Glucose Repression in Saccharomyces cerevisiae. Genetics 108, 845. https://doi.org/10.1093/GENETICS/108.4.845.

Olsen, J. V., Macek, B., Lange, O., Makarov, A., Horning, S., and Mann, M. (2007). Higher-energy C-trap dissociation for peptide modification analysis. Nat. Methods 2007 49 *4*, 709–712. https://doi.org/10.1038/nmeth1060.

Paix, A., Folkmann, A., Rasoloson, D., and Seydoux, G. (2015). High efficiency, homology-directed genome editing in Caenorhabditis elegans using CRISPR-Cas9ribonucleoprotein complexes. Genetics 201, 47–54. https://doi.org/10.1534/genetics.115.179382.

Patro, R., Duggal, G., Love, M.I., Irizarry, R.A., and Kingsford, C. (2017). Salmon provides fast and bias-aware quantification of transcript expression. Nat. Methods 2017 144 *14*, 417–419. https://doi.org/10.1038/nmeth.4197.

Poplin, R., Ruano-Rubio, V., DePristo, M.A., Fennell, T.J., Carneiro, M.O., Auwera, G.A. Van der, Kling, D.E., Gauthier, L.D., Levy-Moonshine, A., Roazen, D., et al. (2018). Scaling accurate genetic variant discovery to tens of thousands of samples. BioRxiv 201178. https://doi.org/10.1101/201178.

Raab, J.R., Resnick, S., and Magnuson, T. (2015). Genome-Wide Transcriptional Regulation Mediated by Biochemically Distinct SWI/SNF Complexes. PLOS Genet. 11, e1005748. https://doi.org/10.1371/JOURNAL.PGEN.1005748.

Ramírez, F., Ryan, D.P., Grüning, B., Bhardwaj, V., Kilpert, F., Richter, A.S., Heyne, S., Dündar, F., and Manke, T. (2016). deepTools2: a next generation web server for deep-sequencing data analysis. Nucleic Acids Res. 44, W160–W165. https://doi.org/10.1093/NAR/GKW257.

Rappsilber, J., Mann, M., and Ishihama, Y. (2007). Protocol for micro-purification, enrichment, pre-fractionation and storage of peptides for proteomics using StageTips. Nat. Protoc. 2007 28 2, 1896–1906. https://doi.org/10.1038/nprot.2007.261.

Reed, B., and Jennings, M. (2011). Guidance on the housing and care of zebrafish Danio rerio.

Riedel, C.G., Dowen, R.H., Lourenco, G.F., Kirienko, N. V., Heimbucher, T., West, J.A., Bowman, S.K., Kingston, R.E., Dillin, A., Asara, J.M., et al. (2013). DAF-16 employs the chromatin remodeller SWI/SNF to promote stress resistance and longevity. Nat. Cell Biol. 2013 155 15, 491–501. https://doi.org/10.1038/ncb2720.

Ruijtenberg, S., and van den Heuvel, S. (2016). Coordinating cell proliferation and differentiation: Antagonism between cell cycle regulators and cell type-specific gene expression. Cell Cycle 15, 196–212. https://doi.org/10.1080/15384101.2015.1120925.

Saha, A., Wittmeyer, J., and Cairns, B.R. (2006). Chromatin remodelling: the industrial revolution of DNA around histones. Nat. Rev. 7, 437–447. https://doi.org/10.1038/nrm1945.

Sawa, H., Kouike, H., and Okano, H. (2000). Components of the SWI/SNF Complex Are Required for Asymmetric Cell Division in C. elegans. Mol. Cell 6, 617–624. .

Schick, S., Rendeiro, A.F., Runggatscher, K., Ringler, A., Boidol, B., Hinkel, M., Májek, P., Vulliard, L., Penz, T., Parapatics, K., et al. (2019). Systematic characterization of BAF mutations provides insights into intracomplex synthetic lethalities in human cancers. Nat. Genet. 51, 1399–1410. https://doi.org/10.1038/S41588-019-0477-9.

Schindelin, J., Arganda-Carreras, I., Frise, E., Kaynig, V., Longair, M., Pietzsch, T., Preibisch, S., Rueden, C., Saalfeld, S., Schmid, B., et al. (2012). Fiji: an open-source platform for biological-image analysis. Nat. Methods 9, 676–682. https://doi.org/10.1038/NMETH.2019.

Serizay, J., Dong, Y., Jänes, J., Chesney, M., Cerrato, C., and Ahringer, J. (2020). Distinctive regulatory architectures of germline-active and somatic genes in C. elegans. Genome Res. 31, 1752– 1765. https://doi.org/10.1101/GR.265934.120/-/DC1.

Sherwood, D.R., and Sternberg, P.W. (2003). Anchor cell invasion into the vulval epithelium in C. elegans. Dev. Cell 5, 21–31. https://doi.org/10.1016/S1534-5807(03)00168-0.

Smith, J.J., Xiao, Y., Parsan, N., Medwig-Kinney, T.N., Martinez, M.A.Q., Moore, F.E.Q., Palmisano, N.J., Kohrman, A.Q., Reyes, M.C.D., Adikes, R.C., et al. (2022). The SWI/SNF chromatin remodeling assemblies BAF and PBAF differentially regulate cell cycle exit and cellular invasion in vivo. PLoS Genet. 18. https://doi.org/10.1371/JOURNAL.PGEN.1009981.

Sokpor, G., Xie, Y., Rosenbusch, J., and Tuoc, T. (2017). Chromatin remodeling BAF (SWI/SNF) complexes in neural development and disorders. Front. Mol. Neurosci. 10, 243. https://doi.org/10.3389/FNMOL.2017.00243/BIBTEX.

Stern, M., Jensen, R., and Herskowitz, I. (1984). Five SWI genes are required for expression of the HO gene in yeast. J. Mol. Biol. 178, 853–868. https://doi.org/10.1016/0022-2836(84)90315-2.

Sulston, J.E., and Horvitz, H.R. (1977). Post-embryonic Cell Lineages of the Nematode, Caenorhabditis elegans. Dev. Biol. 156, 110–156. .

Tyanova, S., Temu, T., Sinitcyn, P., Carlson, A., Hein, M.Y., Geiger, T., Mann, M., and Cox, J. (2016). The Perseus computational platform for comprehensive analysis of (prote)omics data. Nat. Methods 2016 139 13, 731–740. https://doi.org/10.1038/nmeth.3901.

Wang, X., Lee, R.S., Alver, B.H., Haswell, J.R., Wang, S., Mieczkowski, J., Drier, Y., Gillespie, S.M., Archer, T.C., Wu, J.N., et al. (2016). SMARCB1-mediated SWI/SNF complex function is essential for enhancer regulation. Nat. Genet. 2016 492 49, 289–295. https://doi.org/10.1038/ng.3746.

Wang, Y., Argiles-Castillo, D., Kane, E.I., Zhou, A., and Spratt, D.E. (2020). HECT E3 ubiquitin ligases - emerging insights into their biological roles and disease relevance. J. Cell Sci. 133. https://doi.org/10.1242/jcs.228072.

Wilson, B.G., and Roberts, C.W.M. (2011). SWI/SNF nucleosome remodellers and cancer. Nat. Rev. 11, 481–492. https://doi.org/10.1038/nrc3068.

Yang, W., Dierking, K., and Schulenburg, H. (2016). WormExp: a web-based application for a Caenorhabditis elegans-specific gene expression enrichment analysis. Bioinformatics 32, 943–945. https://doi.org/10.1093/BIOINFORMATICS/BTV667.

Zhang, Y., Liu, T., Meyer, C.A., Eeckhoute, J., Johnson, D.S., Bernstein, B.E., Nussbaum, C., Myers, R.M., Brown, M., Li, W., et al. (2008). Model-based analysis of ChIP-Seq (MACS). Genome Biol. 9, 1–9. https://doi.org/10.1186/GB-2008-9-9-R137/FIGURES/3.

Zheng, N., and Shabek, N. (2017). Ubiquitin Ligases: Structure, Function, and Regulation. Annu. Rev. Biochem. 86, 129–157. https://doi.org/10.1146/annurev-biochem-060815-014922.

